# Exploring the genomic and proteomic variations of SARS-CoV-2 spike glycoprotein: a computational biology approach

**DOI:** 10.1101/2020.04.07.030924

**Authors:** Syed Mohammad Lokman, Md. Rasheduzzaman, Asma Salauddin, Rocktim Barua, Afsana Yeasmin Tanzina, Meheadi Hasan Rumi, Md. Imran Hossain, Amam Zonaed Siddiki, Adnan Mannan, Md. Mahbub Hasan

**Affiliations:** Department of Genetic Engineering & Biotechnology, Faculty of Biological Sciences, University of Chittagong, Chattogram-4331, Bangladesh; Department of Pathology and Parasitology, Chittagong Veterinary and Animal Sciences University, Chattogram-4202, Bangladesh

**Keywords:** SARS-CoV-2, Spike protein, Sequence analysis, COVID-19, Genomic variants

## Abstract

The newly identified SARS-CoV-2 has now been reported from around 183 countries with more than a million confirmed human cases including more than 68000 deaths. The genomes of SARS-COV-2 strains isolated from different parts of the world are now available and the unique features of constituent genes and proteins have gotten substantial attention recently. Spike glycoprotein is widely considered as a possible target to be explored because of its role during the entry of coronaviruses into host cells. We analyzed 320 whole-genome sequences and 320 spike protein sequences of SARS-CoV-2 using multiple sequence alignment tools. In this study, 483 unique variations have been identified among the genomes including 25 non-synonymous mutations and one deletion in the spike protein of SARS-CoV-2. Among the 26 variations detected, 12 variations were located at the N-terminal domain and 6 variations at the receptor-binding domain (RBD) which might alter the interaction with receptor molecules. In addition, 22 amino acid insertions were identified in the spike protein of SARS-CoV-2 in comparison with that of SARS-CoV. Phylogenetic analyses of spike protein revealed that Bat coronavirus have a close evolutionary relationship with circulating SARS-CoV-2. The genetic variation analysis data presented in this study can help a better understanding of SARS-CoV-2 pathogenesis. Based on our findings, potential inhibitors can be designed and tested targeting these proposed sites of variation.

## 1. Introduction

Coronavirus disease (COVID-19) is a pandemic manifesting respiratory illness and first reported in Wuhan, Hubei province of China in December 2019. The death toll rose to more than 68,000 among 1,250,000 confirmed cases around the Globe (until April 4, 2020) [1]. The virus causing COVID-19 is named as severe acute respiratory syndrome coronavirus 2 (SARS-CoV-2). Based on the phylogenetic studies, the SARS-CoV-2 is categorized as a member of the genus Betacoronavirus, the same lineage that includes SARS coronavirus (SARS-CoV) [2] that caused SARS (Severe Acute Respiratory Syndrome) in China during 2002 [3]. Recent studies showed that SARS-CoV-2 has a close relationship with bat SARS-like CoVs [4,5], though the intermediate hosts for zoonotic transmission of SARS-CoV-2 from bats to humans remain undiscovered.

SARS-CoV-2 has been identified as an enveloped, single-stranded positive-sense RNA virus with a genome size of approximately 29.9 kb encoding 27 proteins from 14 ORFs including 15 non-structural, 8 accessory and 4 major structural proteins. Two-thirds of the viral RNA harbours the first ORF (ORF1ab) dedicated for translating polyprotein 1a (pp1a) and polyprotein 1ab (pp1ab), which later undergo proteolytic cleavage to form 15 non-structural proteins. Spike glycoprotein (S), membrane (M), envelope (E) and nucleocapsid (N) are the four major structural proteins of SARS-CoV-2 [[6],[7]]. Interestingly, S glycoprotein is characterized as the critical determinant for viral entry into host cells which consists of two functional subunits namely S1 and S2. The S1 subunit recognizes and binds to the host receptor through the receptor-binding domain (RBD) whereas S2 is responsible for fusion with the host cell membrane [[8],[9],[10]]. MERS-CoV uses dipeptidyl peptidase-4 (DPP4) as entry receptor [11] whereas SARS-CoV and SARS-CoV-2 utilize ACE-2 (angiotensin converting enzyme-2) [12], abundantly available in lung alveolar epithelial cells and enterocytes, suggesting S glycoprotein as a potential drug target to halt the entry of SARS-CoV-2 [13].

According to recent reports, neutralizing antibodies are generated in response to the entry and fusion of surface-exposed S protein (mainly RBD domain) which is predicted to be an important target for vaccine candidates [[10],[14],[15]]. However, SARS-CoV-2 has emerged with remarkable properties like glutamine-rich 42 aa long exclusive molecular signature (DSQQTVGQQDGSEDNQTTTIQTIVEVQPQLEMELTPVVQTIE) in position 983-1024 of polyprotein 1ab (pp1ab) [16], diversified receptor-binding domain (RBD), unique furin cleavage site (PRRAR↓SV) at S1/S2 boundary in S glycoprotein which could play roles in viral pathogenesis, diagnosis and treatment [17]. To date, few genomic variations of SARS-CoV-2 are reported [[18],[19]]. There is growing evidence that spike protein, a 1273 amino acid long glycoprotein having multiple domains, possibly plays a major role in SARS-CoV-2 pathogenesis. Viral entry to the host cell is initiated by the receptor-binding domain (RBD) of S1 head. Upon receptor-binding, proteolytic cleavage occurs at S1/S2 cleavage site and two heptad repeats (HR) of S2 stalk form a six-helix bundle structure triggering the release of the fusion peptide. As it comes into close proximity to the transmembrane anchor (TM), the TM domain facilitates membrane destabilization required for fusion between virus-host membranes [[20],[21]]. Insights into the sequence variations of S glycoprotein among available genomes are key to understanding the biology of SARS-CoV-2 infection, developing antiviral treatments and vaccines. In this study, we have analyzed 320 genomic sequences of SARS-CoV-2 to identify mutations between the available genomes followed by the amino acid variations in the glycoprotein S to foresee their impact on the viral entry to host cell from structural biology viewpoint.

## 2. Methods and Materials

### 2.1 Dataset

All available sequences (320 whole genome and surface glycoprotein sequences of SARS-CoV-2) related to the COVID-19 pandemic were retrieved from NCBI Virus Variation Resource repository (https://www.ncbi.nlm.nih.gov/labs/virus/) [22]. In addition, all 40 S glycoprotein sequences from different coronavirus families were retrieved for phylogenetic analysis. The NCBI reference sequence of SARS-CoV-2 S glycoprotein, accession number YP_009724390 was used as the canonical sequence for the analyses of spike protein variants.

### 2.2 Phylogenetic analysis

Variant analyses of SARS-CoV-2 genomes were performed in the Genome Detective Coronavirus Typing Tool Version 1.13 which is specially designed for this virus (https://www.genomedetective.com/app/typingtool/cov/) [23]. For multiple sequence alignment (MSA), Genome Detective Coronavirus Typing Tool uses a reference dataset of 431 whole genome sequences (WGS) where 386 WGS were from known nine coronavirus species. The dataset was then aligned with MUSCLE [24]. Entropy (H(x)) plot of nucleotide variations in SARS-CoV-2 genome was constructed using BioEdit [25]. MEGA X (version 10.1.7) was used to construct the MSAs and the phylogenetic tree using pairwise alignment and neighbor-joining methods in ClustalW [26,27]. Tree structure was validated by running the analysis on 1000 bootstraps [28] replications dataset and the evolutionary distances were calculated using the Poisson correction method [29].

### 2.3 Homology modeling of S glycoprotein

Variant sequences of SARS-CoV-2 were modeled in Swiss-Model [30] using the Cryo-EM spike protein structure of SARS-CoV-2 (PDB ID 6VSB) as a template. The overall quality of models was assessed in RAMPAGE server [31] by generating Ramachandran plots (Supplementary Table 1). PyMol and BIOVIA Discovery Studio were used for structure visualization and superpose [32,33].

## 3. Results

### 3.1 Genomic variations of SARS-CoV-2

Multiple sequence alignment of the available 320 genomes of SARS-CoV-2 were performed and 483 variations were found throughout the 29,903 bp long SARS-CoV-2 genome with in total 115 variations in UTR region, 130 synonymous variations that cause no amino acid alteration, 228 non-synonymous variations causing change in amino acid residue, 16 INDELs, and 2 variations in non-coding region (Supplementary Table 2). Among the 483 variations, 40 variations (14 synonymous, 25 non-synonymous mutations and one deletion) were observed in the region of ORF S that encodes S glycoprotein which is responsible for viral fusion and entry into the host cell [34]. Notable that, most of the SARS-CoV-2 genome sequences were deposited from the USA (250) and China (50) (Supplementary Fig. 1). Positional variability of the SARS-CoV-2 genome was calculated from the MSA of 320 SARS-CoV-2 whole genomes as a measure of Entropy value (H(x)) [35]. Excluding 5′ and 3′ UTR, ten hotspot of hypervariable position were identified, of which seven were located at ORF1ab (1059C>T, 3037C>T, 8782C>T, 14408C>T, 17747C>T, 17858A>G, 18060C>T) and one at ORF S (23403A>G), ORF3a (25563G>T), and ORF8 (28144T>C) respectively. The variability at position 8782 and 28144 were found to be the highest among the other hotspots (Fig. 1).

**Fig. 1:**
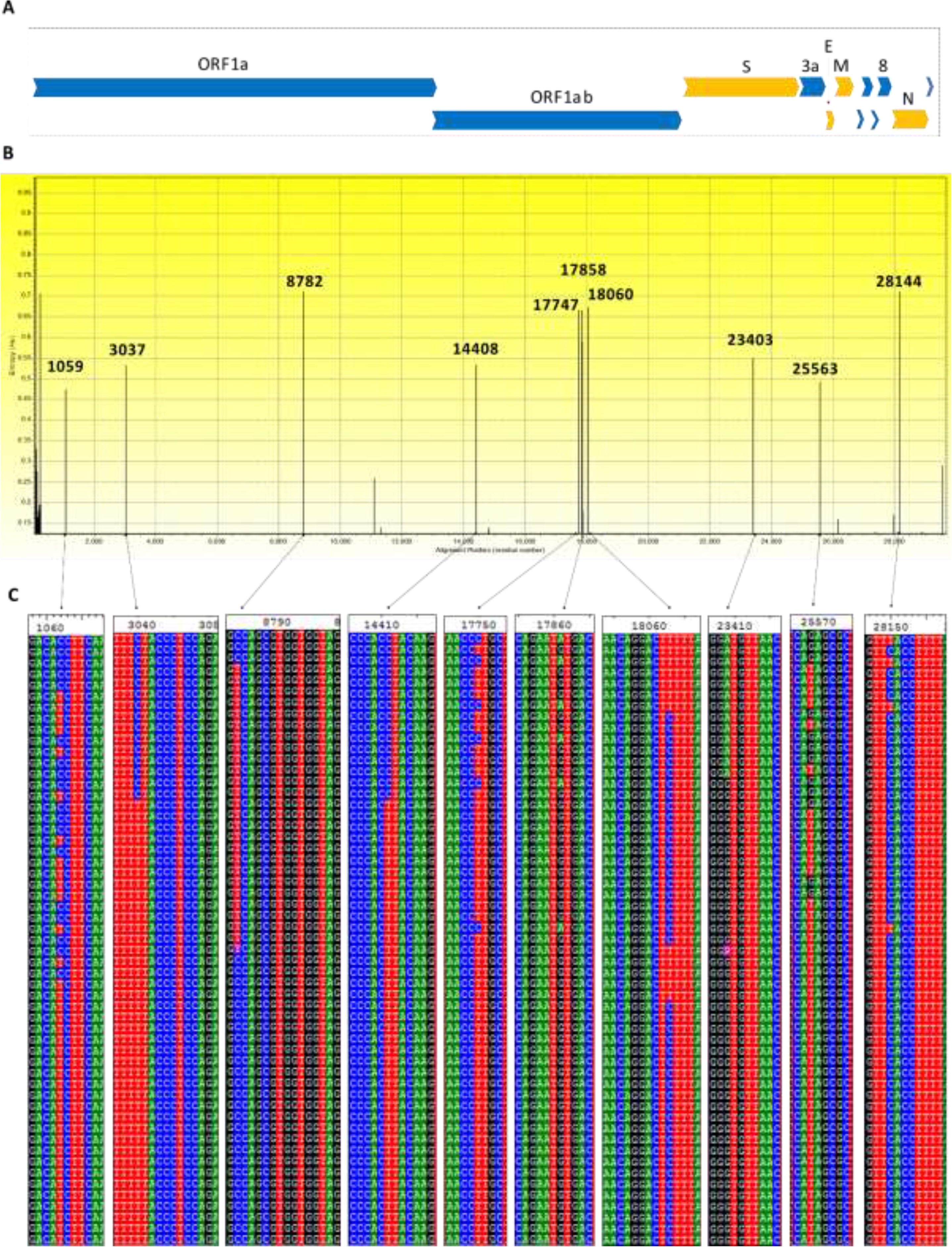
Nucleotide sequence variation among 320 SARS-CoV-2 whole genomes. A. Positional organization of major structural protein-encoding genes in orange color (S = Spike protein, E = Envelope protein, M = Membrane protein, N = Nucleocapsid protein) and accessory protein ORFS in blue colors. B. Variability within 320 SARS-CoV-2 genomic sequences represented by entropy (H(x)) value across genomic location. Two highest frequency of alterations were found at position 8785 of ORF1a and 28144 of ORF8. C. The respective alignment view of each highly variable regions.

### 3.2 Phylogenetic Analysis of S glycoprotein

The phylogenetic analysis of a total of 66 sequences (26 unique SARS-CoV-2 and 40 different coronavirus S glycoprotein sequences) was performed. The evolutionary distances showed that all the SARS-CoV-2 spike proteins cluster in the same node of the phylogenetic tree confirming the sequences are similar to Refseq YP_009724390 (Fig. 2). Bat coronaviruses has a close evolutionary relationship as different strains were found in the nearest outgroups and clades (Bat coronavirus BM48-31, Bat hp-beta coronavirus, Bat coronavirus HKU9) conferring that coronavirus has vast geographical spread and bat is the most prevalent host (Fig. 2). In other clades, the clusters were speculated through different hosts which may describe the evolutionary changes of surface glycoprotein due to cross species transmission. Viral hosts reported from different spots at different times is indicative of possible recombination.

**Fig. 2:**
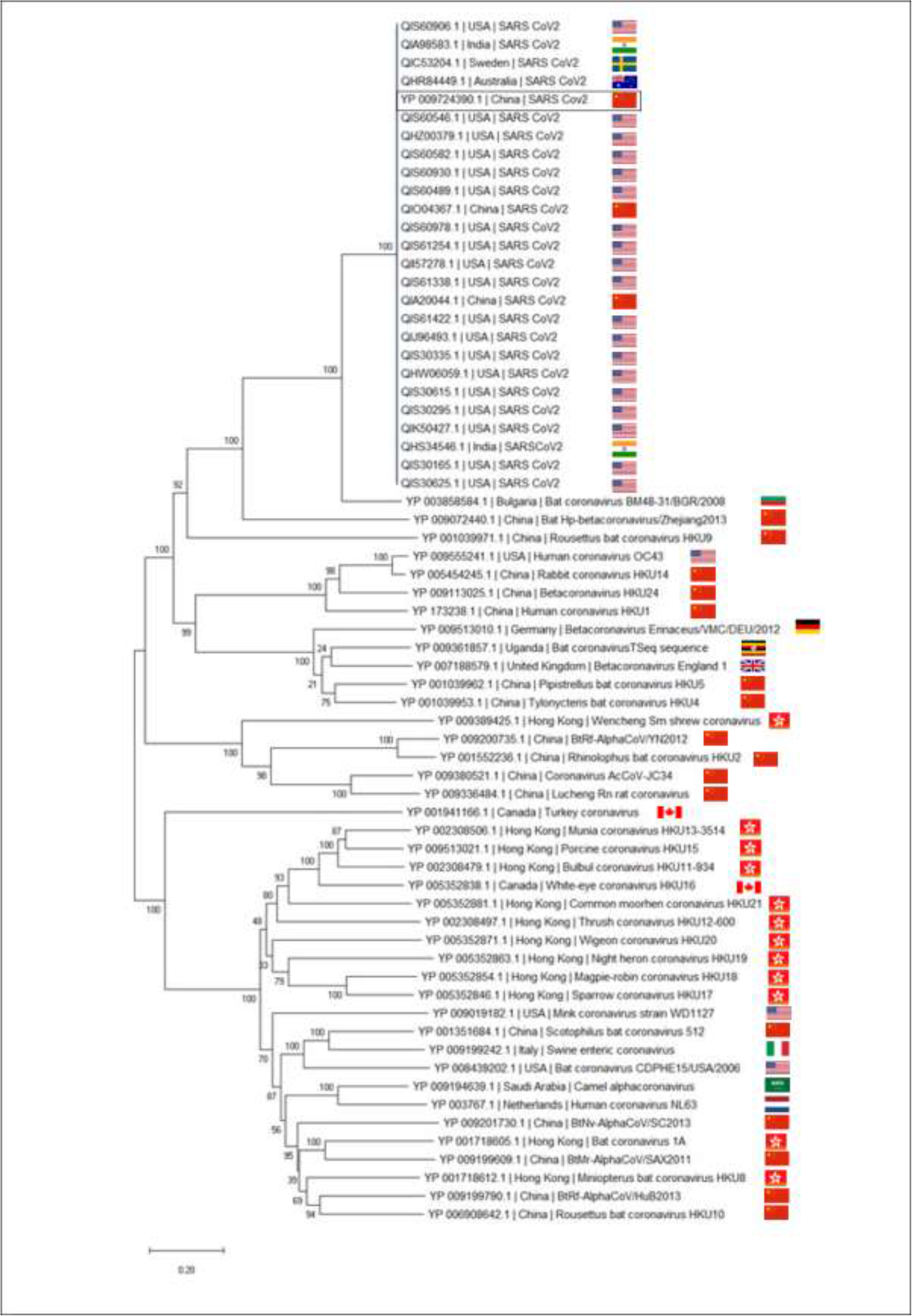
**Sequence phylogeny of SARS-CoV-2 spike glycoprotein variants and other coronavirus spike proteins based on amino acid sequences**, retrieved from NCBI database using neighbor-joining methods in ClustalW and tree structure was validated by running the analysis on 1000 bootstraps. The branch length is indicated in the scale bar. The accession number YP_009724390 represents identical sequences out of SARS-CoV-2 spike proteins.

### 3.3 SARS-CoV-2 Spike Protein Variation Analysis

The S glycoprotein sequences of SARS-CoV-2 were retrieved from the NCBI Virus Variation Resource repository and aligned using ClustalW. The position of SARS-CoV-2 spike protein domains was measured by aligning with the SARS-CoV spike protein (Fig. 3) [36,37]. From the sequence identity matrix, 26 unique variants including the canonical sequence of SARS-CoV-2 spike glycoprotein (YP_009724390) were identified to have 25 substitution and a deletion (Fig. 4A and Supplementary Table 3). 215 sequences were found identical with SARS-CoV-2 S protein reference sequence (YP_009724390) while 64 sequences were identical with the same variation of D614G (Supplementary Table 4, 5). Among all the variations, twelve (Y28N, T29I, H49Y, L54F, N74K, E96D, D111N, Y145Del, F157L, G181V, S221W, and S247R) were located at N-terminal domain (NTD). Another 6 variations (A348T, R408I, G476S, V483A, H519Q, A520S) were found at the receptor-binding domain (RBD) while only two variations (A930V, and D936Y) were found at heptad repeat 1 (HR1) domain. Single variations were found in signal peptide (L5F) domain, sub-domain-2 (D614G), sub-domain-3 (A1078V), heptad repeat 2 domain (D1168H), and cytoplasmic tail domain (D1259H) each. Notable that substitution of Cystine by Phenylalanine was observed at 19 amino acids upstream of the fusion peptide domain (Fig. 4A). The mutation of Aspartic acid to Glycine at position 614 was observed 71 times with entropy value over 0.5 among the available 320 SARS-CoV-2 spike protein sequences (Supplementary Fig. 2).

**Fig. 3:**
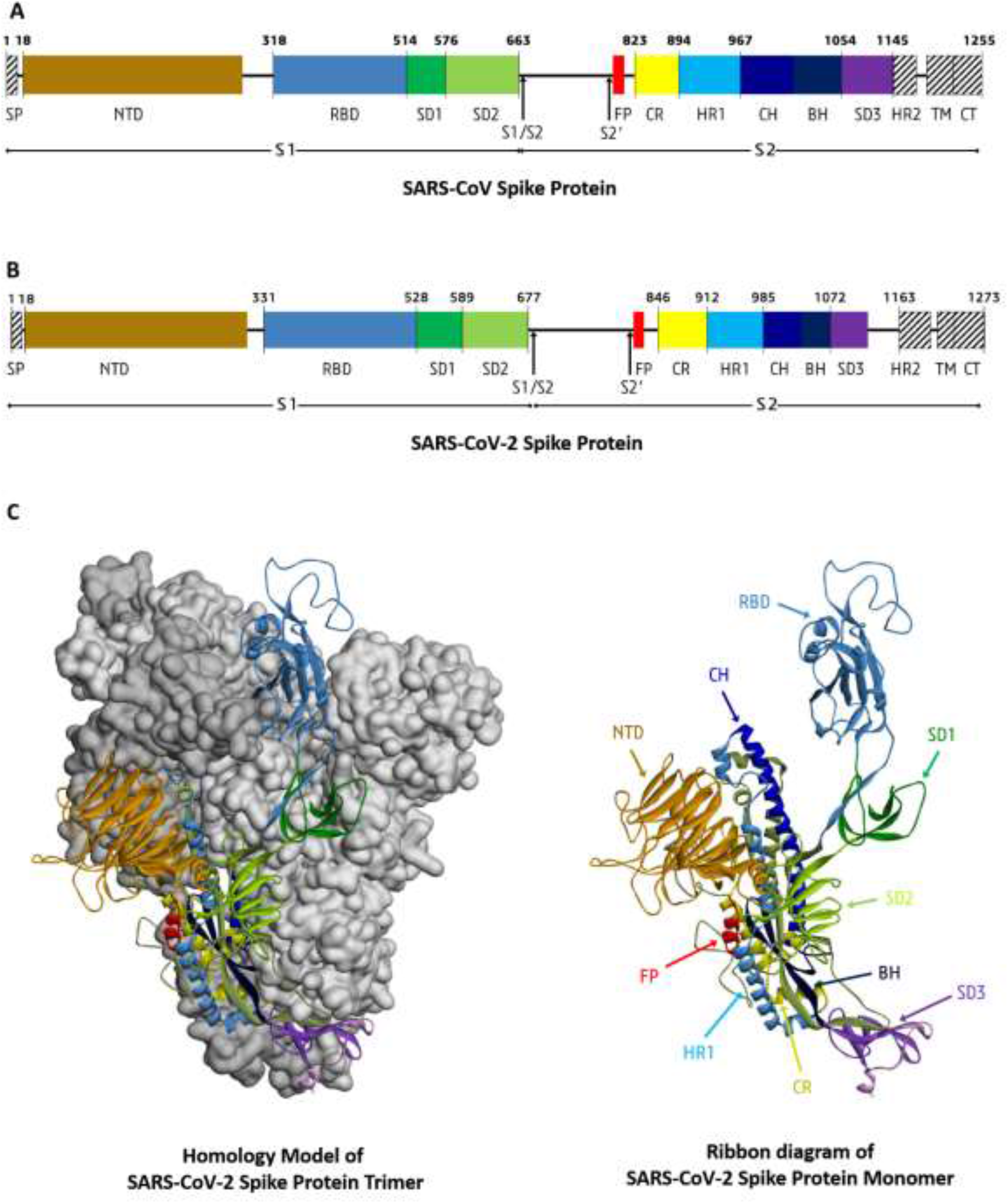
Overall architecture of the SARS-CoV-2 S glycoprotein. A.Schematic diagram of the SARS-CoV S glycoprotein showing domain organization (Reconstructed from Y. Yuan et al., 2017 and M. Gui et al., 2017). B. Schematic domain organization diagram of the SARS-CoV-2 S glycoprotein constructed by aligning with SARS-CoV S protein domain. C. Homology model of SARS-CoV-2 S protein reference sequence YP_009724390 with PDB:6VSB. S protein trimer with two protomers surface shadowed (left). Ribbon diagram of SARS-CoV-2 S glycoprotein monomer from B. Here, NTD: N-terminal domain; RBD: receptor-binding domain; SD: subdomain; CR: connecting region; HR: heptad repeat; CH: central helix; BH: b-hairpin; FP: fusion peptide; TM: transmembrane domain; CT: cytoplasmic tail.

**Fig. 4:**
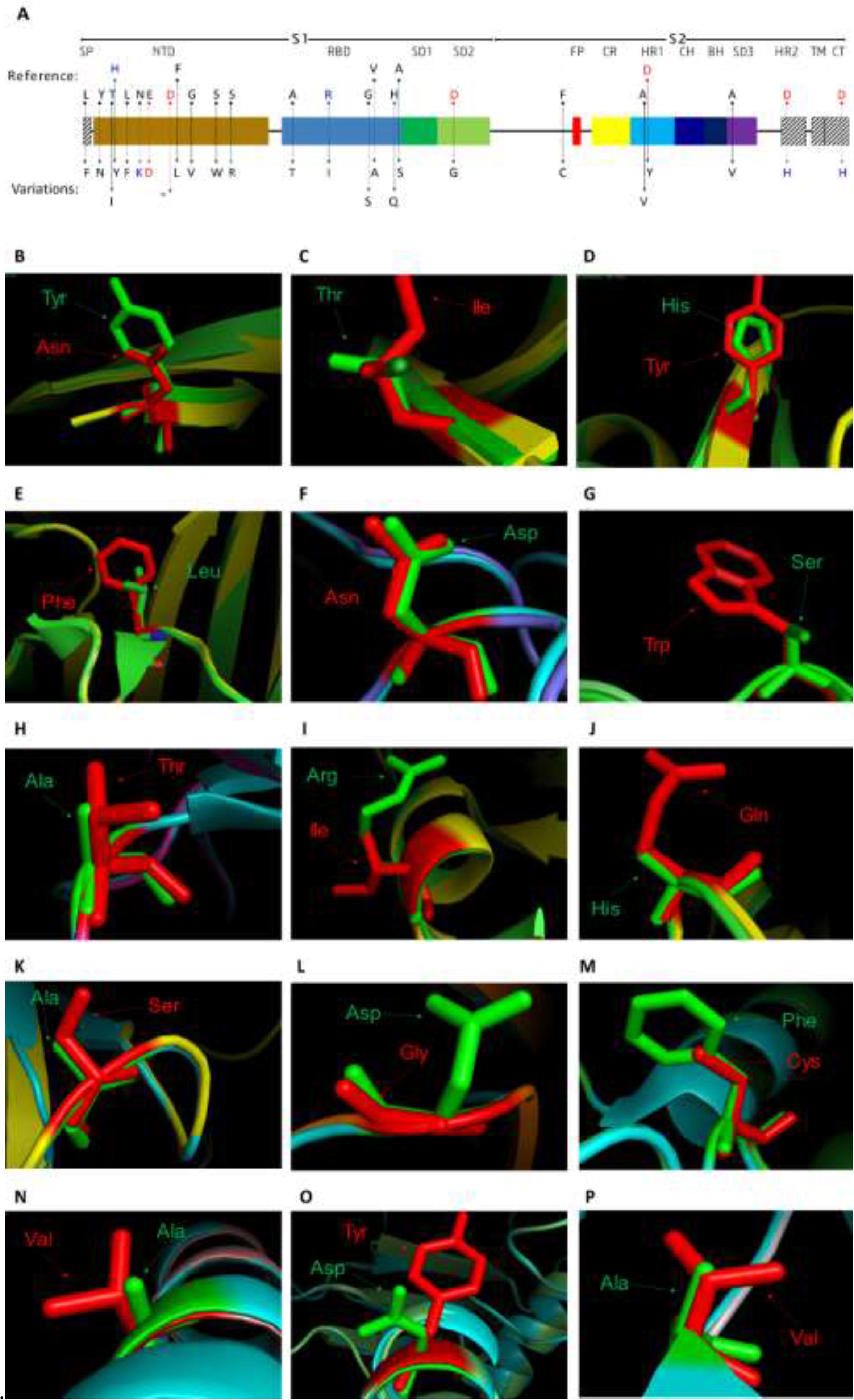
Variability within 320 SARS-CoV-2 S protein sequences. A. Schematic representations of mutations across the spike protein domain organization. Blue, red, and black color represented charge of amino acid residue as positive, negative, and neutral respectively. B-N, Superposed structure of SARS-CoV-2 spike protein variants with the Cryo-EM structure of SARS-CoV-2 Spike Protein (PDB: 6VSB). Template residues are indicated by green color and variants’ residues are indicated as red color. Here, B: Y28N, C: T29I, D: H49Y, E: L54F, F: D111N, G: S221W, H: A348T, I: R408I, J: H519Q, K: A520S, L: D614G, M: F797C, N: A930V, O: D936Y, and P: A1078V.

Alterations of amino acid residual charge from positive to neutral (H49Y, R408I, H519Q), negative to neutral (D111N, D614G, D936Y), negative to positive (D1168H, D1259H), and neutral to positive (N74K, S247R) were seen in variants QHW06059, QHS34546, QIS61422, QIS61338, QIK50427, QIS30615, QIS60978, QIS60582, QIO04367, and QHR84449 respectively due to substitution of amino acid that differ in charge. The remaining 15 variants were mutated with the amino acids that are similar in charge (Fig. 4 A). The SARS-CoV-2 spike protein variants were superposed with the cryo-electron microscopic structure of SARS-CoV-2 spike protein [8]. L5F, N74K, E96D, F157L, G181V, S247R, G476S, V483A, D1168H, and D1259H variants were excluded from superposition due to absence of respective residues in the 3D structure of template (PDB: 6VSB). The superposition showed that most of the residual change were causing incorporation of bulky amino acid residues (T29I, H49Y, L54F, S221W, A348T, H519Q, A520S, A930V, D936Y, and A1078V) in place of smaller size residue except Y28N, D111N, R408I, D614G, and F797C (Fig. 4 B-P).

The sequence comparison of spike glycoprotein between SARS-CoV-2 variants and SARS-CoV (Uniprot ID: P59594) revealed nearly 77.46% similarity and identified the presence of additional 22 amino acids in SARS-CoV-2 spike protein variants resulting from a total of 5 insertions (Supplementary Fig. 3). Among these, the major insertion consisting of 7 amino acids (GTNGTKR) at position 72-78 followed by 4 amino acids (NKSW) at position 149-152 and 6 amino acids (SYLTPG) at 247-252 occurred in N-terminal domain. Insertion of Glycine at 482 was found in receptor binding domain, preceding another insertion of 4 amino acids (NSPR) at position 679-682, just upstream of S1/S2 cleavage site that leads to form a furin-like cleavage site (PRRARS) in the S protein variants of SARS-CoV-2 (Supplementary Fig. 3). The S2 subunit of spike protein, especially the heptad repeat region 2, fusion peptide domain, transmembrane domain, and cytoplasmic tail were found to be highly conserved in the SARS-CoV and the SARS-CoV-2 variants while the S1 subunit was more diverse, specifically the N-terminal domain (NTD) and receptor-binding domain (RBD).

## 4. Discussion

COVID 19 is one of the most contagious pandemics the world has ever had with 1,250,000 confirmed cases to date (April 4, 2020) and the cases have increased as high as 5 times in less than a month [1]. Phylogenetic analysis showed that the SARS-CoV-2 is a unique coronavirus presumably related to Bat coronavirus (BM48-31, Hp-betacoronavirus). During this study, we investigated the available genomes of SARS-CoV-2 and found variations in 483 positions resulting in 130 synonymous and 228 non-synonymous variants. Out of them, 25 non-synonymous variants were observed in the spike protein of SARS-CoV-2. Viral spike protein is thought to have a crucial role in drug and vaccine development as reported previously in managing the viruses like SARS-CoV and MERS-CoV [[15],[38],[39],[40]]. Likewise, a number of studies targeting SARS-CoV-2 spike protein have been undertaken for the therapeutic measures [41], but the unique structural and functional details of SARS-CoV-2 spike protein are still under scrutiny. We also found a variant (R408I) at receptor binding domain (RBD) that mutated from positively charged Arginine residue to neutral and smaller sized Isoleucine residue (Fig. 4 I). This change might alter the interaction of viral RBD with the host receptor because the R408 residue of SARS-CoV-2 is known to interact with the ACE2 receptor for viral entry [42]. Similarly, alterations of RBD (G476S, V483A, H519Q, and A520S) also could affect the interaction of SARS-CoV-2 spike protein with other molecules which require further investigations. QIA98583 and QIS30615 variants were found to have an alteration of Alanine to Valine (A930V), and Aspartic acid to Tyrosine (D936Y) respectively in the alpha helix of the HR1 domain. Previous reports have indicated that HR1 domain plays a significant role in viral fusion and entry by forming helical bundles with HR2, and mutations including alanine substitution by valine (A1168V) in HR1 region are predominantly responsible for conferring resistance to mouse hepatitis coronaviruses against HR2 derived peptide entry inhibitors [43]. This study hypothesizes the mutation (A930V) found in that of SARS-CoV-2 might also have a role in the emergence of drug-resistance virus strains. Also, the mutation (D1168H) found in the heptad repeat 2 (HR) SARS-CoV-2 could play a vital role in viral pathogenesis.

The SARS-CoV-2 S protein contains additional furin protease cleavage site, PRRARS, in S1/S2 domain which is conserved among all 320 sequences as revealed during this study (Supplementary Fig. 3). This unique signature is thought to make the SARS-CoV-2 more virulent than SARS-CoV and regarded as novel features of the viral pathogenesis (ref 11). According to previous reports the more the host cell protease can process the coronavirus S can accelerate viral tropism accordingly in influenza virus [[9],[44],[45],[46]]. Apart from that, this could also promote viruses to escape antiviral therapies targeting transmembrane protease TMPRSS2 (ClinicalTrials.gov, NCT04321096) which is well reported protease to cleave at S1/S2 of S glycoprotein [47]. Comparative analyses between SARS-CoV and SARS-CoV-2 spike glycoprotein showed 77% similarity between them where the most diverse region was the N-terminal domain and receptor binding domain. The insertion of 17 additional amino acid residues in the N-terminal domain of SARS-CoV-2 and its high sequence diversity suggests that it may have a role in binding with other cell receptors in humans. This is because the N-terminal domain could function as the receptor-binding domain of various coronaviruses [[34],[36]]. Similar phenomenon has been observed in mouse hepatitis coronavirus (MHV) and porcine transmissible gastroenteritis coronavirus (TGEV) where the N-terminal domain is reported to be attached with the host entry receptor [48,49]. The variation analyses in amino acids indicated the structural features of different domains of the SARS-CoV-2 spike proteins. However, to identify the actual role of involvement of S glycoprotein, a larger dataset regarding genomics and proteomics of SARS-CoV-2 is required as this protein is vital to understand the viral pathogenicity, evolution and development of therapeutics. Further analyses of all the S glycoprotein and SARS-CoV-2 genomes with different epidemiological aspects is warranted to get the pathogenesis of SARS-CoV-2.

