## Supplementary file for "Exploring the genomic and proteomic variations of SARS-CoV-2 spike glycoprotein: a computational biology approach"

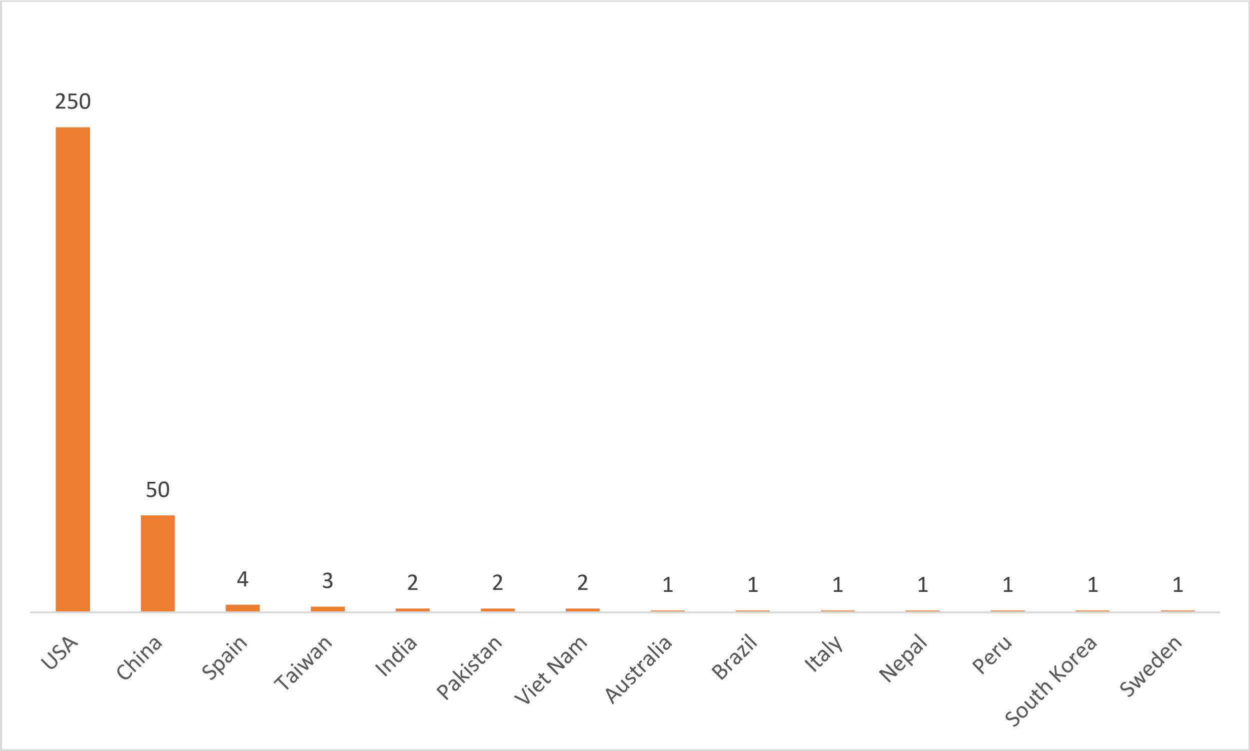


Supplementary Fig. 1. **Genomic data sources based on geo location of 320 SARS-CoV-2 whole genome sequences.**

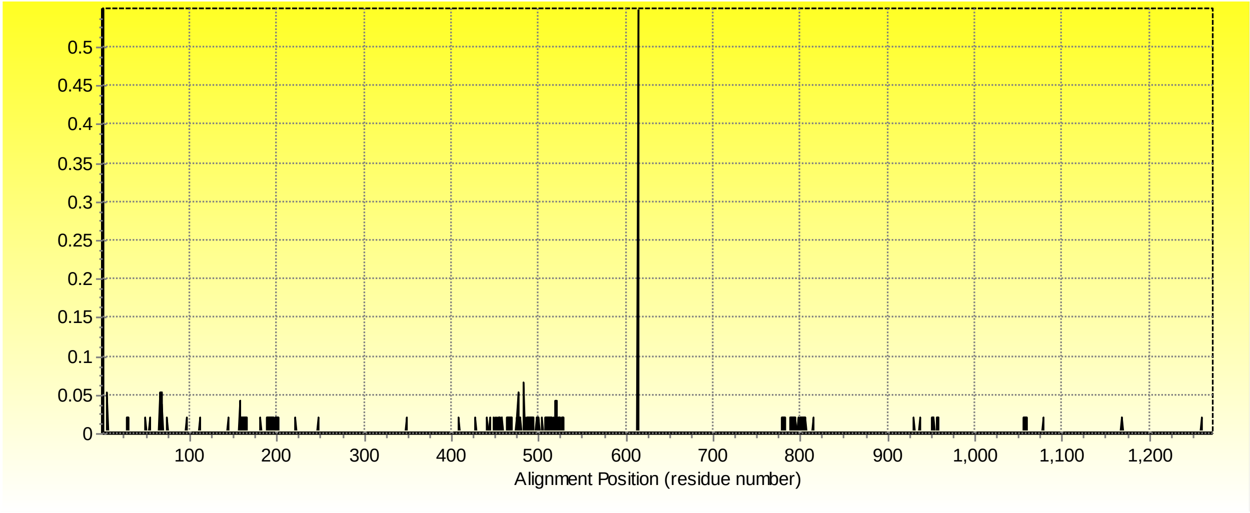


Supplementary Fig. 2. **Entropy plot measuring positional variability of SARS-CoV-2 spike protein sequences.**

```
YP_009724390  1    MFVFLVLLPL VSSQCVN--L TTRTQLPPAY TN--SFTRGV YYPDKVFRSS VLHSTQDLFL
QHS34546.1    1    MFVFLVLLPL VSSQCVN--L TTRTQLPPAY TN--SFTRGV YYPDKVFRSS VLHSTQDLFL
QIS60906.1    1    MFVFFVLLPL VSSQCVN--L TTRTQLPPAY TN--SFTRGV YYPDKVFRSS VLHSTQDLFL
QIS60930.1    1    MFVFLVLLPL VSSQCVN--L TTRTQLPPAY TN--SFTRGV YYPDKVFRSS VLHSTQDLFL
QIS61338.1    1    MFVFLVLLPL VSSQCVN--L TTRTQLPPAY TN--SFTRGV YYPDKVFRSS VLHSTQDLFL
QIS60489.1    1    MFVFLVLLPL VSSQCVN--L TTRTQLPPAY TN--SFTRGV YYPDKVFRSS VLHSTQDLFL
QIS30625.1    1    MFVFLVLLPL VSSQCVN--L TTRTQLPPAY TN--SFTRGV YYPDKVFRSS VLHSTQDLFL
QIS30335.1    1    MFVFLVLLPL VSSQCVN--L TTRTQLPPAY TN--SFTRGV YYPDKVFRSS VLHSTQDLFL
QIC53204.1    1    MFVFLVLLPL VSSQCVN--L TTRTQLPPAY TN--SFTRGV YYPDKVFRSS VLHSTQDLFL
QHW06059.1    1    MFVFLVLLPL VSSQCVN--L TTRTQLPPAY TN--SFTRGV YYPDKVFRSS VLYSTQDLFL
QHZ00379.1    1    MFVFLVLLPL VSSQCVN--L TTRTQLPPAY TN--SFTRGV YYPDKVFRSS VLHSTQDLFL
QHR84449.1    1    MFVFLVLLPL VSSQCVN--L TTRTQLPPAY TN--SFTRGV YYPDKVFRSS VLHSTQDLFL
QIJ96493.1    1    MFVFLVLLPL VSSQCVN--L TTRTQLPPAY TN--SFTRGV YYPDKVFRSS VLHSTQDLFL
QIA98583.1    1    MFVFLVLLPL VSSQCVN--L TTRTQLPPAY TN--SFTRGV YYPDKVFRSS VLHSTQDLFL
QII57278.1    1    MFVFLVLLPL VSSQCVN--L TTRTQLPPAY TN--SFTRGV YYPDKVFRSS VLHSTQDLFL
QIA20044.1    1    MFVFLVLLPL VSSQCVN--L TTRTQLPPAN TN--SFTRGV YYPDKVFRSS VLHSTQDLFL
QIS60546.1    1    MFVFLVLLPL VSSQCVN--L TTRTQLPPAY IN--SFTRGV YYPDKVFRSS VLHSTQDLFL
QIS60582.1    1    MFVFLVLLPL VSSQCVN--L TTRTQLPPAY TN--SFTRGV YYPDKVFRSS VLHSTQDLFL
QIS61254.1    1    MFVFLVLLPL VSSQCVN--L TTRTQLPPAY TN--SFTRGV YYPDKVFRSS VLHSTQDLFL
QIS61422.1    1    MFVFLVLLPL VSSQCVN--L TTRTQLPPAY TN--SFTRGV YYPDKVFRSS VLHSTQDLFL
QIS60978.1    1    MFVFLVLLPL VSSQCVN--L TTRTQLPPAY TN--SFTRGV YYPDKVFRSS VLHSTQDLFL
QIO04367.1    1    MFVFLVLLPL VSSQCVN--L TTRTQLPPAY TN--SFTRGV YYPDKVFRSS VLHSTQDLFL
QIS30295.1    1    MFVFLVLLPL VSSQCVN--L TTRTQLPPAY TN--SFTRGV YYPDKVFRSS VLHSTQDFFL
QIS30615.1    1    MFVFLVLLPL VSSQCVN--L TTRTQLPPAY TN--SFTRGV YYPDKVFRSS VLHSTQDLFL
QIK50427.1    1    MFVFLVLLPL VSSQCVN--L TTRTQLPPAY TN--SFTRGV YYPDKVFRSS VLHSTQDLFL
QIS30165.1    1    MFVFLVLLPL VSSQCVN--L TTRTQLPPAY TN--SFTRGV YYPDKVFRSS VLHSTQDLFL
SARS-CoV      1    MFIFLLFLTL TSGSDLDRCT TFDDVQAPNY TQHTSSMRGV YYPDEIFRSD TLYLTQDLFL
                   ________________     __________________________________________________
                          SP                                  NTD

YP_009724390  57   PFFSNVTWFH AIHVSGTNGT KRFDNPVLPF NDGVYFASTE KSNIIRGWIF GTTLDSKTQS
QHS34546.1    57   PFFSNVTWFH AIHVSGTNGT KRFDNPVLPF NDGVYFASTE KSNIIRGWIF GTTLDSKTQS
QIS60906.1    57   PFFSNVTWFH AIHVSGTNGT KRFDNPVLPF NDGVYFASTE KSNIIRGWIF GTTLDSKTQS
QIS60930.1    57   PFFSNVTWFH AIHVSGTNGT KRFDNPVLPF NDGVYFASTD KSNIIRGWIF GTTLDSKTQS
QIS61338.1    57   PFFSNVTWFH AIHVSGTNGT KRFDNPVLPF NDGVYFASTE KSNIIRGWIF GTTLNSKTQS
QIS60489.1    57   PFFSNVTWFH AIHVSGTNGT KRFDNPVLPF NDGVYFASTE KSNIIRGWIF GTTLDSKTQS
QIS30625.1    57   PFFSNVTWFH AIHVSGTNGT KRFDNPVLPF NDGVYFASTE KSNIIRGWIF GTTLDSKTQS
QIS30335.1    57   PFFSNVTWFH AIHVSGTNGT KRFDNPVLPF NDGVYFASTE KSNIIRGWIF GTTLDSKTQS
QIC53204.1    57   PFFSNVTWFH AIHVSGTNGT KRFDNPVLPF NDGVYFASTE KSNIIRGWIF GTTLDSKTQS
QHW06059.1    57   PFFSNVTWFH AIHVSGTNGT KRFDNPVLPF NDGVYFASTE KSNIIRGWIF GTTLDSKTQS
QHZ00379.1    57   PFFSNVTWFH AIHVSGTNGT KRFDNPVLPF NDGVYFASTE KSNIIRGWIF GTTLDSKTQS
QHR84449.1    57   PFFSNVTWFH AIHVSGTNGT KRFDNPVLPF NDGVYFASTE KSNIIRGWIF GTTLDSKTQS
QIJ96493.1    57   PFFSNVTWFH AIHVSGTNGT KRFDNPVLPF NDGVYFASTE KSNIIRGWIF GTTLDSKTQS
QIA98583.1    57   PFFSNVTWFH AIHVSGTNGT KRFDNPVLPF NDGVYFASTE KSNIIRGWIF GTTLDSKTQS
QII57278.1    57   PFFSNVTWFH AIHVSGTNGT KRFDNPVLPF NDGVYFASTE KSNIIRGWIF GTTLDSKTQS
QIA20044.1    57   PFFSNVTWFH AIHVSGTNGT KRFDNPVLPF NDGVYFASTE KSNIIRGWIF GTTLDSKTQS
QIS60546.1    57   PFFSNVTWFH AIHVSGTNGT KRFDNPVLPF NDGVYFASTE KSNIIRGWIF GTTLDSKTQS
QIS60582.1    57   PFFSNVTWFH AIHVSGTNGT KRFDNPVLPF NDGVYFASTE KSNIIRGWIF GTTLDSKTQS
QIS61254.1    57   PFFSNVTWFH AIHVSGTNGT KRFDNPVLPF NDGVYFASTE KSNIIRGWIF GTTLDSKTQS
QIS61422.1    57   PFFSNVTWFH AIHVSGTNGT KRFDNPVLPF NDGVYFASTE KSNIIRGWIF GTTLDSKTQS
QIS60978.1    57   PFFSNVTWFH AIHVSGTNGT KRFDNPVLPF NDGVYFASTE KSNIIRGWIF GTTLDSKTQS
QIO04367.1    57   PFFSNVTWFH AIHVSGTKGT KRFDNPVLPF NDGVYFASTE KSNIIRGWIF GTTLDSKTQS
QIS30295.1    57   PFFSNVTWFH AIHVSGTNGT KRFDNPVLPF NDGVYFASTE KSNIIRGWIF GTTLDSKTQS
QIS30615.1    57   PFFSNVTWFH AIHVSGTNGT KRFDNPVLPF NDGVYFASTE KSNIIRGWIF GTTLDSKTQS
QIK50427.1    57   PFFSNVTWFH AIHVSGTNGT KRFDNPVLPF NDGVYFASTE KSNIIRGWIF GTTLDSKTQS
QIS30165.1    57   PFFSNVTWFH AIHVSGTNGT KRFDNPVLPF NDGVYFASTE KSNIIRGWIF GTTLDSKTQS
SARS-CoV      61   PFYSNVTGFH TINHT----- --FGNPVIPF KDGIYFAATE KSNVVRGWVF GSTMNNKSQS
```

NTD

```
YP_009724390   117  LLIVNNATNV VIKVCEFQFC NDPFLGVYYH KNNKSWMESE FRVYSSANNC TFEYVSQPFL
QHS34546.1     117  LLIVNNATNV VIKVCEFQFC NDPFLGV-YH KNNKSWMESE FRVYSSANNC TFEYVSQPFL
QIS60906.1     117  LLIVNNATNV VIKVCEFQFC NDPFLGVYYH KNNKSWMESE FRVYSSANNC TFEYVSQPFL
QIS60930.1     117  LLIVNNATNV VIKVCEFQFC NDPFLGVYYH KNNKSWMESE FRVYSSANNC TFEYVSQPFL
QIS61338.1     117  LLIVNNATNV VIKVCEFQFC NDPFLGVYYH KNNKSWMESE FRVYSSANNC TFEYVSQPFL
QIS60489.1     117  LLIVNNATNV VIKVCEFQFC NDPFLGVYYH KNNKSWMESE FRVYSSANNC TFEYVSQPFL
QIS30625.1     117  LLIVNNATNV VIKVCEFQFC NDPFLGVYYH KNNKSWMESE FRVYSSANNC TFEYVSQPFL
QIS30335.1     117  LLIVNNATNV VIKVCEFQFC NDPFLGVYYH KNNKSWMESE FRVYSSANNC TFEYVSQPFL
QIC53204.1     117  LLIVNNATNV VIKVCEFQFC NDPFLGVYYH KNNKSWMESE FRVYSSANNC TFEYVSQPFL
QHW06059.1     117  LLIVNNATNV VIKVCEFQFC NDPFLGVYYH KNNKSWMESE FRVYSSANNC TFEYVSQPFL
QHZ00379.1     117  LLIVNNATNV VIKVCEFQFC NDPFLGVYYH KNNKSWMESE FRVYSSANNC TFEYVSQPFL
QHR84449.1     117  LLIVNNATNV VIKVCEFQFC NDPFLGVYYH KNNKSWMESE FRVYSSANNC TFEYVSQPFL
QIJ96493.1     117  LLIVNNATNV VIKVCEFQFC NDPFLGVYYH KNNKSWMESE FRVYSSANNC TFEYVSQPFL
QIA98583.1     117  LLIVNNATNV VIKVCEFQFC NDPFLGVYYH KNNKSWMESE FRVYSSANNC TFEYVSQPFL
QII57278.1     117  LLIVNNATNV VIKVCEFQFC NDPFLGVYYH KNNKSWMESE LRVYSSANNC TFEYVSQPFL
QIA20044.1     117  LLIVNNATNV VIKVCEFQFC NDPFLGVYYH KNNKSWMESE FRVYSSANNC TFEYVSQPFL
QIS60546.1     117  LLIVNNATNV VIKVCEFQFC NDPFLGVYYH KNNKSWMESE FRVYSSANNC TFEYVSQPFL
QIS60582.1     117  LLIVNNATNV VIKVCEFQFC NDPFLGVYYH KNNKSWMESE FRVYSSANNC TFEYVSQPFL
QIS61254.1     117  LLIVNNATNV VIKVCEFQFC NDPFLGVYYH KNNKSWMESE FRVYSSANNC TFEYVSQPFL
QIS61422.1     117  LLIVNNATNV VIKVCEFQFC NDPFLGVYYH KNNKSWMESE FRVYSSANNC TFEYVSQPFL
QIS60978.1     117  LLIVNNATNV VIKVCEFQFC NDPFLGVYYH KNNKSWMESE FRVYSSANNC TFEYVSQPFL
QIO04367.1     117  LLIVNNATNV VIKVCEFQFC NDPFLGVYYH KNNKSWMESE FRVYSSANNC TFEYVSQPFL
QIS30295.1     117  LLIVNNATNV VIKVCEFQFC NDPFLGVYYH KNNKSWMESE FRVYSSANNC TFEYVSQPFL
QIS30615.1     117  LLIVNNATNV VIKVCEFQFC NDPFLGVYYH KNNKSWMESE FRVYSSANNC TFEYVSQPFL
QIK50427.1     117  LLIVNNATNV VIKVCEFQFC NDPFLGVYYH KNNKSWMESE FRVYSSANNC TFEYVSQPFL
QIS30165.1     117  LLIVNNATNV VIKVCEFQFC NDPFLGVYYH KNNKSWMESE FRVYSSANNC TFEYVSQPFL
SARS-CoV       114  VIIINNSTNV VIRACNFELC DNPFFAVSKP MG----TQTH TMIFDNAFNC TFEYISDAFS
```

NTD

```
YP_009724390   177  MDLEGKQGNF KNLREFVFKN IDGYFKIYSK HTPINLVRDL PQGFSALEPL VDLPIGINIT
QHS34546.1     176  MDLEGKQGNF KNLREFVFKN IDGYFKIYSK HTPINLVRDL PQGFSALEPL VDLPIGINIT
QIS60906.1     177  MDLEGKQGNF KNLREFVFKN IDGYFKIYSK HTPINLVRDL PQGFSALEPL VDLPIGINIT
QIS60930.1     177  MDLEGKQGNF KNLREFVFKN IDGYFKIYSK HTPINLVRDL PQGFSALEPL VDLPIGINIT
QIS61338.1     177  MDLEGKQGNF KNLREFVFKN IDGYFKIYSK HTPINLVRDL PQGFSALEPL VDLPIGINIT
QIS60489.1     177  MDLEGKQGNF KNLREFVFKN IDGYFKIYSK HTPINLVRDL PQGFSALEPL VDLPIGINIT
QIS30625.1     177  MDLEGKQGNF KNLREFVFKN IDGYFKIYSK HTPINLVRDL PQGFSALEPL VDLPIGINIT
QIS30335.1     177  MDLEGKQGNF KNLREFVFKN IDGYFKIYSK HTPINLVRDL PQGFSALEPL VDLPIGINIT
QIC53204.1     177  MDLEGKQGNF KNLREFVFKN IDGYFKIYSK HTPINLVRDL PQGFSALEPL VDLPIGINIT
QHW06059.1     177  MDLEGKQGNF KNLREFVFKN IDGYFKIYSK HTPINLVRDL PQGFSALEPL VDLPIGINIT
QHZ00379.1     177  MDLEGKQGNF KNLREFVFKN IDGYFKIYSK HTPINLVRDL PQGFWALEPL VDLPIGINIT
QHR84449.1     177  MDLEGKQGNF KNLREFVFKN IDGYFKIYSK HTPINLVRDL PQGFSALEPL VDLPIGINIT
QIJ96493.1     177  MDLEVKQGNF KNLREFVFKN IDGYFKIYSK HTPINLVRDL PQGFSALEPL VDLPIGINIT
QIA98583.1     177  MDLEGKQGNF KNLREFVFKN IDGYFKIYSK HTPINLVRDL PQGFSALEPL VDLPIGINIT
QII57278.1     177  MDLEGKQGNF KNLREFVFKN IDGYFKIYSK HTPINLVRDL PQGFSALEPL VDLPIGINIT
QIA20044.1     177  MDLEGKQGNF KNLREFVFKN IDGYFKIYSK HTPINLVRDL PQGFSALEPL VDLPIGINIT
QIS60546.1     177  MDLEGKQGNF KNLREFVFKN IDGYFKIYSK HTPINLVRDL PQGFSALEPL VDLPIGINIT
QIS60582.1     177  MDLEGKQGNF KNLREFVFKN IDGYFKIYSK HTPINLVRDL PQGFSALEPL VDLPIGINIT
QIS61254.1     177  MDLEGKQGNF KNLREFVFKN IDGYFKIYSK HTPINLVRDL PQGFSALEPL VDLPIGINIT
QIS61422.1     177  MDLEGKQGNF KNLREFVFKN IDGYFKIYSK HTPINLVRDL PQGFSALEPL VDLPIGINIT
QIS60978.1     177  MDLEGKQGNF KNLREFVFKN IDGYFKIYSK HTPINLVRDL PQGFSALEPL VDLPIGINIT
QIO04367.1     177  MDLEGKQGNF KNLREFVFKN IDGYFKIYSK HTPINLVRDL PQGFSALEPL VDLPIGINIT
QIS30295.1     177  MDLEGKQGNF KNLREFVFKN IDGYFKIYSK HTPINLVRDL PQGFSALEPL VDLPIGINIT
QIS30615.1     177  MDLEGKQGNF KNLREFVFKN IDGYFKIYSK HTPINLVRDL PQGFSALEPL VDLPIGINIT
QIK50427.1     177  MDLEGKQGNF KNLREFVFKN IDGYFKIYSK HTPINLVRDL PQGFSALEPL VDLPIGINIT
QIS30165.1     177  MDLEGKQGNF KNLREFVFKN IDGYFKIYSK HTPINLVRDL PQGFSALEPL VDLPIGINIT
SARS-CoV       170  LDVSEKSGNF KHLREFVFKN KDGFLYVYKG YQPIDVVRDL PSGFNTLKPI FKLPLGINIT
```

NTD

| | | | | | | | |
|---|---|---|---|---|---|---|---|
| YP_009724390 | 237 | RFQTLLALHR | SYLTPGDSSS | GWTAGAAAYY | VGYLQPRTFL | LKYNENGTIT | DAVDCALDPL |
| QHS34546.1 | 236 | RFQTLLALHR | SYLTPGDSSS | GWTAGAAAYY | VGYLQPRTFL | LKYNENGTIT | DAVDCALDPL |
| QIS60906.1 | 237 | RFQTLLALHR | SYLTPGDSSS | GWTAGAAAYY | VGYLQPRTFL | LKYNENGTIT | DAVDCALDPL |
| QIS60930.1 | 237 | RFQTLLALHR | SYLTPGDSSS | GWTAGAAAYY | VGYLQPRTFL | LKYNENGTIT | DAVDCALDPL |
| QIS61338.1 | 237 | RFQTLLALHR | SYLTPGDSSS | GWTAGAAAYY | VGYLQPRTFL | LKYNENGTIT | DAVDCALDPL |
| QIS60489.1 | 237 | RFQTLLALHR | SYLTPGDSSS | GWTAGAAAYY | VGYLQPRTFL | LKYNENGTIT | DAVDCALDPL |
| QIS30625.1 | 237 | RFQTLLALHR | SYLTPGDSSS | GWTAGAAAYY | VGYLQPRTFL | LKYNENGTIT | DAVDCALDPL |
| QIS30335.1 | 237 | RFQTLLALHR | SYLTPGDSSS | GWTAGAAAYY | VGYLQPRTFL | LKYNENGTIT | DAVDCALDPL |
| QIC53204.1 | 237 | RFQTLLALHR | SYLTPGDSSS | GWTAGAAAYY | VGYLQPRTFL | LKYNENGTIT | DAVDCALDPL |
| QHW06059.1 | 237 | RFQTLLALHR | SYLTPGDSSS | GWTAGAAAYY | VGYLQPRTFL | LKYNENGTIT | DAVDCALDPL |
| QHZ00379.1 | 237 | RFQTLLALHR | SYLTPGDSSS | GWTAGAAAYY | VGYLQPRTFL | LKYNENGTIT | DAVDCALDPL |
| QHR84449.1 | 237 | RFQTLLALHR | RYLTPGDSSS | GWTAGAAAYY | VGYLQPRTFL | LKYNENGTIT | DAVDCALDPL |
| QIJ96493.1 | 237 | RFQTLLALHR | SYLTPGDSSS | GWTAGAAAYY | VGYLQPRTFL | LKYNENGTIT | DAVDCALDPL |
| QIA98583.1 | 237 | RFQTLLALHR | SYLTPGDSSS | GWTAGAAAYY | VGYLQPRTFL | LKYNENGTIT | DAVDCALDPL |
| QII57278.1 | 237 | RFQTLLALHR | SYLTPGDSSS | GWTAGAAAYY | VGYLQPRTFL | LKYNENGTIT | DAVDCALDPL |
| QIA20044.1 | 237 | RFQTLLALHR | SYLTPGDSSS | GWTAGAAAYY | VGYLQPRTFL | LKYNENGTIT | DAVDCALDPL |
| QIS60546.1 | 237 | RFQTLLALHR | SYLTPGDSSS | GWTAGAAAYY | VGYLQPRTFL | LKYNENGTIT | DAVDCALDPL |
| QIS60582.1 | 237 | RFQTLLALHR | SYLTPGDSSS | GWTAGAAAYY | VGYLQPRTFL | LKYNENGTIT | DAVDCALDPL |
| QIS61254.1 | 237 | RFQTLLALHR | SYLTPGDSSS | GWTAGAAAYY | VGYLQPRTFL | LKYNENGTIT | DAVDCALDPL |
| QIS61422.1 | 237 | RFQTLLALHR | SYLTPGDSSS | GWTAGAAAYY | VGYLQPRTFL | LKYNENGTIT | DAVDCALDPL |
| QIS60978.1 | 237 | RFQTLLALHR | SYLTPGDSSS | GWTAGAAAYY | VGYLQPRTFL | LKYNENGTIT | DAVDCALDPL |
| QIO04367.1 | 237 | RFQTLLALHR | SYLTPGDSSS | GWTAGAAAYY | VGYLQPRTFL | LKYNENGTIT | DAVDCALDPL |
| QIS30295.1 | 237 | RFQTLLALHR | SYLTPGDSSS | GWTAGAAAYY | VGYLQPRTFL | LKYNENGTIT | DAVDCALDPL |
| QIS30615.1 | 237 | RFQTLLALHR | SYLTPGDSSS | GWTAGAAAYY | VGYLQPRTFL | LKYNENGTIT | DAVDCALDPL |
| QIK50427.1 | 237 | RFQTLLALHR | SYLTPGDSSS | GWTAGAAAYY | VGYLQPRTFL | LKYNENGTIT | DAVDCALDPL |
| QIS30165.1 | 237 | RFQTLLALHR | SYLTPGDSSS | GWTAGAAAYY | VGYLQPRTFL | LKYNENGTIT | DAVDCALDPL |
| SARS-CoV | 230 | NFRAILTAFS | ------PAQD | IWGTSAAAYF | VGYLKPTTFM | LKYDENGTIT | DAVDCSQNPL |

NTD

| | | | | | | | |
|---|---|---|---|---|---|---|---|
| YP_009724390 | 297 | SETKCTLKSF | TVEKGIYQTS | NFRVQPTESI | VRFPNITNLC | PFGEVFNATR | FASVYAWNRK |
| QHS34546.1 | 296 | SETKCTLKSF | TVEKGIYQTS | NFRVQPTESI | VRFPNITNLC | PFGEVFNATR | FASVYAWNRK |
| QIS60906.1 | 297 | SETKCTLKSF | TVEKGIYQTS | NFRVQPTESI | VRFPNITNLC | PFGEVFNATR | FASVYAWNRK |
| QIS60930.1 | 297 | SETKCTLKSF | TVEKGIYQTS | NFRVQPTESI | VRFPNITNLC | PFGEVFNATR | FASVYAWNRK |
| QIS61338.1 | 297 | SETKCTLKSF | TVEKGIYQTS | NFRVQPTESI | VRFPNITNLC | PFGEVFNATR | FASVYAWNRK |
| QIS60489.1 | 297 | SETKCTLKSF | TVEKGIYQTS | NFRVQPTESI | VRFPNITNLC | PFGEVFNATR | FASVYAWNRK |
| QIS30625.1 | 297 | SETKCTLKSF | TVEKGIYQTS | NFRVQPTESI | VRFPNITNLC | PFGEVFNATR | FASVYAWNRK |
| QIS30335.1 | 297 | SETKCTLKSF | TVEKGIYQTS | NFRVQPTESI | VRFPNITNLC | PFGEVFNATR | FTSVYAWNRK |
| QIC53204.1 | 297 | SETKCTLKSF | TVEKGIYQTS | NFRVQPTESI | VRFPNITNLC | PFGEVFNATR | FASVYAWNRK |
| QHW06059.1 | 297 | SETKCTLKSF | TVEKGIYQTS | NFRVQPTESI | VRFPNITNLC | PFGEVFNATR | FASVYAWNRK |
| QHZ00379.1 | 297 | SETKCTLKSF | TVEKGIYQTS | NFRVQPTESI | VRFPNITNLC | PFGEVFNATR | FASVYAWNRK |
| QHR84449.1 | 297 | SETKCTLKSF | TVEKGIYQTS | NFRVQPTESI | VRFPNITNLC | PFGEVFNATR | FASVYAWNRK |
| QIJ96493.1 | 297 | SETKCTLKSF | TVEKGIYQTS | NFRVQPTESI | VRFPNITNLC | PFGEVFNATR | FASVYAWNRK |
| QIA98583.1 | 297 | SETKCTLKSF | TVEKGIYQTS | NFRVQPTESI | VRFPNITNLC | PFGEVFNATR | FASVYAWNRK |
| QII57278.1 | 297 | SETKCTLKSF | TVEKGIYQTS | NFRVQPTESI | VRFPNITNLC | PFGEVFNATR | FASVYAWNRK |
| QIA20044.1 | 297 | SETKCTLKSF | TVEKGIYQTS | NFRVQPTESI | VRFPNITNLC | PFGEVFNATR | FASVYAWNRK |
| QIS60546.1 | 297 | SETKCTLKSF | TVEKGIYQTS | NFRVQPTESI | VRFPNITNLC | PFGEVFNATR | FASVYAWNRK |
| QIS60582.1 | 297 | SETKCTLKSF | TVEKGIYQTS | NFRVQPTESI | VRFPNITNLC | PFGEVFNATR | FASVYAWNRK |
| QIS61254.1 | 297 | SETKCTLKSF | TVEKGIYQTS | NFRVQPTESI | VRFPNITNLC | PFGEVFNATR | FASVYAWNRK |
| QIS61422.1 | 297 | SETKCTLKSF | TVEKGIYQTS | NFRVQPTESI | VRFPNITNLC | PFGEVFNATR | FASVYAWNRK |
| QIS60978.1 | 297 | SETKCTLKSF | TVEKGIYQTS | NFRVQPTESI | VRFPNITNLC | PFGEVFNATR | FASVYAWNRK |
| QIO04367.1 | 297 | SETKCTLKSF | TVEKGIYQTS | NFRVQPTESI | VRFPNITNLC | PFGEVFNATR | FASVYAWNRK |
| QIS30295.1 | 297 | SETKCTLKSF | TVEKGIYQTS | NFRVQPTESI | VRFPNITNLC | PFGEVFNATR | FASVYAWNRK |
| QIS30615.1 | 297 | SETKCTLKSF | TVEKGIYQTS | NFRVQPTESI | VRFPNITNLC | PFGEVFNATR | FASVYAWNRK |
| QIK50427.1 | 297 | SETKCTLKSF | TVEKGIYQTS | NFRVQPTESI | VRFPNITNLC | PFGEVFNATR | FASVYAWNRK |
| QIS30165.1 | 297 | SETKCTLKSF | TVEKGIYQTS | NFRVQPTESI | VRFPNITNLC | PFGEVFNATR | FASVYAWNRK |
| SARS-CoV | 284 | AELKCSVKSF | EIDKGIYQTS | NFRVVPSGDV | VRFPNITNLC | PFGEVFNATK | FPSVYAWERK |

NTD RBD

| | | | | | | | |
|---|---|---|---|---|---|---|---|
| YP_009724390 | 357 | RISNCVADYS | VLYNSASFST | FKCYGVSPTK | LNDLCFTNVY | ADSFVIRGDE | VRQIAPGQTG |
| QHS34546.1 | 356 | RISNCVADYS | VLYNSASFST | FKCYGVSPTK | LNDLCFTNVY | ADSFVIRGDE | VIQIAPGQTG |
| QIS60906.1 | 357 | RISNCVADYS | VLYNSASFST | FKCYGVSPTK | LNDLCFTNVY | ADSFVIRGDE | VRQIAPGQTG |
| QIS60930.1 | 357 | RISNCVADYS | VLYNSASFST | FKCYGVSPTK | LNDLCFTNVY | ADSFVIRGDE | VRQIAPGQTG |
| QIS61338.1 | 357 | RISNCVADYS | VLYNSASFST | FKCYGVSPTK | LNDLCFTNVY | ADSFVIRGDE | VRQIAPGQTG |
| QIS60489.1 | 357 | RISNCVADYS | VLYNSASFST | FKCYGVSPTK | LNDLCFTNVY | ADSFVIRGDE | VRQIAPGQTG |
| QIS30625.1 | 357 | RISNCVADYS | VLYNSASFST | FKCYGVSPTK | LNDLCFTNVY | ADSFVIRGDE | VRQIAPGQTG |
| QIS30335.1 | 357 | RISNCVADYS | VLYNSASFST | FKCYGVSPTK | LNDLCFTNVY | ADSFVIRGDE | VRQIAPGQTG |
| QIC53204.1 | 357 | RISNCVADYS | VLYNSASFST | FKCYGVSPTK | LNDLCFTNVY | ADSFVIRGDE | VRQIAPGQTG |
| QHW06059.1 | 357 | RISNCVADYS | VLYNSASFST | FKCYGVSPTK | LNDLCFTNVY | ADSFVIRGDE | VRQIAPGQTG |
| QHZ00379.1 | 357 | RISNCVADYS | VLYNSASFST | FKCYGVSPTK | LNDLCFTNVY | ADSFVIRGDE | VRQIAPGQTG |
| QHR84449.1 | 357 | RISNCVADYS | VLYNSASFST | FKCYGVSPTK | LNDLCFTNVY | ADSFVIRGDE | VRQIAPGQTG |
| QIJ96493.1 | 357 | RISNCVADYS | VLYNSASFST | FKCYGVSPTK | LNDLCFTNVY | ADSFVIRGDE | VRQIAPGQTG |
| QIA98583.1 | 357 | RISNCVADYS | VLYNSASFST | FKCYGVSPTK | LNDLCFTNVY | ADSFVIRGDE | VRQIAPGQTG |
| QII57278.1 | 357 | RISNCVADYS | VLYNSASFST | FKCYGVSPTK | LNDLCFTNVY | ADSFVIRGDE | VRQIAPGQTG |
| QIA20044.1 | 357 | RISNCVADYS | VLYNSASFST | FKCYGVSPTK | LNDLCFTNVY | ADSFVIRGDE | VRQIAPGQTG |
| QIS60546.1 | 357 | RISNCVADYS | VLYNSASFST | FKCYGVSPTK | LNDLCFTNVY | ADSFVIRGDE | VRQIAPGQTG |
| QIS60582.1 | 357 | RISNCVADYS | VLYNSASFST | FKCYGVSPTK | LNDLCFTNVY | ADSFVIRGDE | VRQIAPGQTG |
| QIS61254.1 | 357 | RISNCVADYS | VLYNSASFST | FKCYGVSPTK | LNDLCFTNVY | ADSFVIRGDE | VRQIAPGQTG |
| QIS61422.1 | 357 | RISNCVADYS | VLYNSASFST | FKCYGVSPTK | LNDLCFTNVY | ADSFVIRGDE | VRQIAPGQTG |
| QIS60978.1 | 357 | RISNCVADYS | VLYNSASFST | FKCYGVSPTK | LNDLCFTNVY | ADSFVIRGDE | VRQIAPGQTG |
| QIO04367.1 | 357 | RISNCVADYS | VLYNSASFST | FKCYGVSPTK | LNDLCFTNVY | ADSFVIRGDE | VRQIAPGQTG |
| QIS30295.1 | 357 | RISNCVADYS | VLYNSASFST | FKCYGVSPTK | LNDLCFTNVY | ADSFVIRGDE | VRQIAPGQTG |
| QIS30615.1 | 357 | RISNCVADYS | VLYNSASFST | FKCYGVSPTK | LNDLCFTNVY | ADSFVIRGDE | VRQIAPGQTG |
| QIK50427.1 | 357 | RISNCVADYS | VLYNSASFST | FKCYGVSPTK | LNDLCFTNVY | ADSFVIRGDE | VRQIAPGQTG |
| QIS30165.1 | 357 | RISNCVADYS | VLYNSASFST | FKCYGVSPTK | LNDLCFTNVY | ADSFVIRGDE | VRQIAPGQTG |
| SARS-CoV | 344 | KISNCVADYS | VLYNSTFFST | FKCYGVSATK | LNDLCFSNVY | ADSFVVKGDD | VRQIAPGQTG |

RBD

| | | | | | | | |
|---|---|---|---|---|---|---|---|
| YP_009724390 | 417 | KIADYNYKLP | DDFTGCVIAW | NSNNLDSKVG | GNYNYLYRLF | RKSNLKPFER | DISTEIYQAG |
| QHS34546.1 | 416 | KIADYNYKLP | DDFTGCVIAW | NSNNLDSKVG | GNYNYLYRLF | RKSNLKPFER | DISTEIYQAG |
| QIS60906.1 | 417 | KIADYNYKLP | DDFTGCVIAW | NSNNLDSKVG | GNYNYLYRLF | RKSNLKPFER | DISTEIYQAG |
| QIS60930.1 | 417 | KIADYNYKLP | DDFTGCVIAW | NSNNLDSKVG | GNYNYLYRLF | RKSNLKPFER | DISTEIYQAG |
| QIS61338.1 | 417 | KIADYNYKLP | DDFTGCVIAW | NSNNLDSKVG | GNYNYLYRLF | RKSNLKPFER | DISTEIYQAG |
| QIS60489.1 | 417 | KIADYNYKLP | DDFTGCVIAW | NSNNLDSKVG | GNYNYLYRLF | RKSNLKPFER | DISTEIYQAG |
| QIS30625.1 | 417 | KIADYNYKLP | DDFTGCVIAW | NSNNLDSKVG | GNYNYLYRLF | RKSNLKPFER | DISTEIYQAS |
| QIS30335.1 | 417 | KIADYNYKLP | DDFTGCVIAW | NSNNLDSKVG | GNYNYLYRLF | RKSNLKPFER | DISTEIYQAG |
| QIC53204.1 | 417 | KIADYNYKLP | DDFTGCVIAW | NSNNLDSKVG | GNYNYLYRLF | RKSNLKPFER | DISTEIYQAG |
| QHW06059.1 | 417 | KIADYNYKLP | DDFTGCVIAW | NSNNLDSKVG | GNYNYLYRLF | RKSNLKPFER | DISTEIYQAG |
| QHZ00379.1 | 417 | KIADYNYKLP | DDFTGCVIAW | NSNNLDSKVG | GNYNYLYRLF | RKSNLKPFER | DISTEIYQAG |
| QHR84449.1 | 417 | KIADYNYKLP | DDFTGCVIAW | NSNNLDSKVG | GNYNYLYRLF | RKSNLKPFER | DISTEIYQAG |
| QIJ96493.1 | 417 | KIADYNYKLP | DDFTGCVIAW | NSNNLDSKVG | GNYNYLYRLF | RKSNLKPFER | DISTEIYQAG |
| QIA98583.1 | 417 | KIADYNYKLP | DDFTGCVIAW | NSNNLDSKVG | GNYNYLYRLF | RKSNLKPFER | DISTEIYQAG |
| QII57278.1 | 417 | KIADYNYKLP | DDFTGCVIAW | NSNNLDSKVG | GNYNYLYRLF | RKSNLKPFER | DISTEIYQAG |
| QIA20044.1 | 417 | KIADYNYKLP | DDFTGCVIAW | NSNNLDSKVG | GNYNYLYRLF | RKSNLKPFER | DISTEIYQAG |
| QIS60546.1 | 417 | KIADYNYKLP | DDFTGCVIAW | NSNNLDSKVG | GNYNYLYRLF | RKSNLKPFER | DISTEIYQAG |
| QIS60582.1 | 417 | KIADYNYKLP | DDFTGCVIAW | NSNNLDSKVG | GNYNYLYRLF | RKSNLKPFER | DISTEIYQAG |
| QIS61254.1 | 417 | KIADYNYKLP | DDFTGCVIAW | NSNNLDSKVG | GNYNYLYRLF | RKSNLKPFER | DISTEIYQAG |
| QIS61422.1 | 417 | KIADYNYKLP | DDFTGCVIAW | NSNNLDSKVG | GNYNYLYRLF | RKSNLKPFER | DISTEIYQAG |
| QIS60978.1 | 417 | KIADYNYKLP | DDFTGCVIAW | NSNNLDSKVG | GNYNYLYRLF | RKSNLKPFER | DISTEIYQAG |
| QIO04367.1 | 417 | KIADYNYKLP | DDFTGCVIAW | NSNNLDSKVG | GNYNYLYRLF | RKSNLKPFER | DISTEIYQAG |
| QIS30295.1 | 417 | KIADYNYKLP | DDFTGCVIAW | NSNNLDSKVG | GNYNYLYRLF | RKSNLKPFER | DISTEIYQAG |
| QIS30615.1 | 417 | KIADYNYKLP | DDFTGCVIAW | NSNNLDSKVG | GNYNYLYRLF | RKSNLKPFER | DISTEIYQAG |
| QIK50427.1 | 417 | KIADYNYKLP | DDFTGCVIAW | NSNNLDSKVG | GNYNYLYRLF | RKSNLKPFER | DISTEIYQAG |
| QIS30165.1 | 417 | KIADYNYKLP | DDFTGCVIAW | NSNNLDSKVG | GNYNYLYRLF | RKSNLKPFER | DISTEIYQAG |
| SARS-CoV | 404 | VIADYNYKLP | DDFMGCVLAW | NTRNIDATST | GNYNYKYRYL | RHGKLRPFER | DISNVPFSPD |

RBD

```
YP_009724390   477  STPCNGVEGF NCYFPLQSYG FQPTNGVGYQ PYRVVVLSFE LLHAPATVCG PKKSTNLVKN
QHS34546.1     476  STPCNGVEGF NCYFPLQSYG FQPTNGVGYQ PYRVVVLSFE LLHAPATVCG PKKSTNLVKN
QIS60906.1     477  STPCNGVEGF NCYFPLQSYG FQPTNGVGYQ PYRVVVLSFE LLHAPATVCG PKKSTNLVKN
QIS60930.1     477  STPCNGVEGF NCYFPLQSYG FQPTNGVGYQ PYRVVVLSFE LLHAPATVCG PKKSTNLVKN
QIS61338.1     477  STPCNGVEGF NCYFPLQSYG FQPTNGVGYQ PYRVVVLSFE LLHAPATVCG PKKSTNLVKN
QIS60489.1     477  STPCNGVEGF NCYFPLQSYG FQPTNGVGYQ PYRVVVLSFE LLHSPATVCG PKKSTNLVKN
QIS30625.1     477  STPCNGVEGF NCYFPLQSYG FQPTNGVGYQ PYRVVVLSFE LLHAPATVCG PKKSTNLVKN
QIS30335.1     477  STPCNGVEGF NCYFPLQSYG FQPTNGVGYQ PYRVVVLSFE LLHAPATVCG PKKSTNLVKN
QIC53204.1     477  STPCNGVEGF NCYFPLQSYG FQPTNGVGYQ PYRVVVLSFE LLHAPATVCG PKKSTNLVKN
QHW06059.1     477  STPCNGVEGF NCYFPLQSYG FQPTNGVGYQ PYRVVVLSFE LLHAPATVCG PKKSTNLVKN
QHZ00379.1     477  STPCNGVEGF NCYFPLQSYG FQPTNGVGYQ PYRVVVLSFE LLHAPATVCG PKKSTNLVKN
QHR84449.1     477  STPCNGVEGF NCYFPLQSYG FQPTNGVGYQ PYRVVVLSFE LLHAPATVCG PKKSTNLVKN
QIJ96493.1     477  STPCNGVEGF NCYFPLQSYG FQPTNGVGYQ PYRVVVLSFE LLHAPATVCG PKKSTNLVKN
QIA98583.1     477  STPCNGVEGF NCYFPLQSYG FQPTNGVGYQ PYRVVVLSFE LLHAPATVCG PKKSTNLVKN
QII57278.1     477  STPCNGVEGF NCYFPLQSYG FQPTNGVGYQ PYRVVVLSFE LLHAPATVCG PKKSTNLVKN
QIA20044.1     477  STPCNGVEGF NCYFPLQSYG FQPTNGVGYQ PYRVVVLSFE LLHAPATVCG PKKSTNLVKN
QIS60546.1     477  STPCNGVEGF NCYFPLQSYG FQPTNGVGYQ PYRVVVLSFE LLHAPATVCG PKKSTNLVKN
QIS60582.1     477  STPCNGVEGF NCYFPLQSYG FQPTNGVGYQ PYRVVVLSFE LLHAPATVCG PKKSTNLVKN
QIS61254.1     477  STPCNGVEGF NCYFPLQSYG FQPTNGVGYQ PYRVVVLSFE LLHAPATVCG PKKSTNLVKN
QIS61422.1     477  STPCNGVEGF NCYFPLQSYG FQPTNGVGYQ PYRVVVLSFE LLQAPATVCG PKKSTNLVKN
QIS60978.1     477  STPCNGVEGF NCYFPLQSYG FQPTNGVGYQ PYRVVVLSFE LLHAPATVCG PKKSTNLVKN
QIO04367.1     477  STPCNGVEGF NCYFPLQSYG FQPTNGVGYQ PYRVVVLSFE LLHAPATVCG PKKSTNLVKN
QIS30295.1     477  STPCNGVEGF NCYFPLQSYG FQPTNGVGYQ PYRVVVLSFE LLHAPATVCG PKKSTNLVKN
QIS30615.1     477  STPCNGVEGF NCYFPLQSYG FQPTNGVGYQ PYRVVVLSFE LLHAPATVCG PKKSTNLVKN
QIK50427.1     477  STPCNGVEGF NCYFPLQSYG FQPTNGVGYQ PYRVVVLSFE LLHAPATVCG PKKSTNLVKN
QIS30165.1     477  STPCNGAEGF NCYFPLQSYG FQPTNGVGYQ PYRVVVLSFE LLHAPATVCG PKKSTNLVKN
SARS-CoV       464  GKPCT-PPAL NCYWPLNDYG FYTTTGIGYQ PYRVVVLSFE LLNAPATVCG PKLSTDLIKN
                    ______________________________RBD__________________________ ___SD1____

YP_009724390   537  KCVNFNFNGL TGTGVLTESN KKFLPFQQFG RDIADTTDAV RDPQTLEILD ITPCSFGGVS
QHS34546.1     536  KCVNFNFNGL TGTGVLTESN KKFLPFQQFG RDIADTTDAV RDPQTLEILD ITPCSFGGVS
QIS60906.1     537  KCVNFNFNGL TGTGVLTESN KKFLPFQQFG RDIADTTDAV RDPQTLEILD ITPCSFGGVS
QIS60930.1     537  KCVNFNFNGL TGTGVLTESN KKFLPFQQFG RDIADTTDAV RDPQTLEILD ITPCSFGGVS
QIS61338.1     537  KCVNFNFNGL TGTGVLTESN KKFLPFQQFG RDIADTTDAV RDPQTLEILD ITPCSFGGVS
QIS60489.1     537  KCVNFNFNGL TGTGVLTESN KKFLPFQQFG RDIADTTDAV RDPQTLEILD ITPCSFGGVS
QIS30625.1     537  KCVNFNFNGL TGTGVLTESN KKFLPFQQFG RDIADTTDAV RDPQTLEILD ITPCSFGGVS
QIS30335.1     537  KCVNFNFNGL TGTGVLTESN KKFLPFQQFG RDIADTTDAV RDPQTLEILD ITPCSFGGVS
QIC53204.1     537  KCVNFNFNGL TGTGVLTESN KKFLPFQQFG RDIADTTDAV RDPQTLEILD ITPCSFGGVS
QHW06059.1     537  KCVNFNFNGL TGTGVLTESN KKFLPFQQFG RDIADTTDAV RDPQTLEILD ITPCSFGGVS
QHZ00379.1     537  KCVNFNFNGL TGTGVLTESN KKFLPFQQFG RDIADTTDAV RDPQTLEILD ITPCSFGGVS
QHR84449.1     537  KCVNFNFNGL TGTGVLTESN KKFLPFQQFG RDIADTTDAV RDPQTLEILD ITPCSFGGVS
QIJ96493.1     537  KCVNFNFNGL TGTGVLTESN KKFLPFQQFG RDIADTTDAV RDPQTLEILD ITPCSFGGVS
QIA98583.1     537  KCVNFNFNGL TGTGVLTESN KKFLPFQQFG RDIADTTDAV RDPQTLEILD ITPCSFGGVS
QII57278.1     537  KCVNFNFNGL TGTGVLTESN KKFLPFQQFG RDIADTTDAV RDPQTLEILD ITPCSFGGVS
QIA20044.1     537  KCVNFNFNGL TGTGVLTESN KKFLPFQQFG RDIADTTDAV RDPQTLEILD ITPCSFGGVS
QIS60546.1     537  KCVNFNFNGL TGTGVLTESN KKFLPFQQFG RDIADTTDAV RDPQTLEILD ITPCSFGGVS
QIS60582.1     537  KCVNFNFNGL TGTGVLTESN KKFLPFQQFG RDIADTTDAV RDPQTLEILD ITPCSFGGVS
QIS61254.1     537  KCVNFNFNGL TGTGVLTESN KKFLPFQQFG RDIADTTDAV RDPQTLEILD ITPCSFGGVS
QIS61422.1     537  KCVNFNFNGL TGTGVLTESN KKFLPFQQFG RDIADTTDAV RDPQTLEILD ITPCSFGGVS
QIS60978.1     537  KCVNFNFNGL TGTGVLTESN KKFLPFQQFG RDIADTTDAV RDPQTLEILD ITPCSFGGVS
QIO04367.1     537  KCVNFNFNGL TGTGVLTESN KKFLPFQQFG RDIADTTDAV RDPQTLEILD ITPCSFGGVS
QIS30295.1     537  KCVNFNFNGL TGTGVLTESN KKFLPFQQFG RDIADTTDAV RDPQTLEILD ITPCSFGGVS
QIS30615.1     537  KCVNFNFNGL TGTGVLTESN KKFLPFQQFG RDIADTTDAV RDPQTLEILD ITPCSFGGVS
QIK50427.1     537  KCVNFNFNGL TGTGVLTESN KKFLPFQQFG RDIADTTDAV RDPQTLEILD ITPCSFGGVS
QIS30165.1     537  KCVNFNFNGL TGTGVLTESN KKFLPFQQFG RDIADTTDAV RDPQTLEILD ITPCSFGGVS
SARS-CoV       523  QCVNFNFNGL TGTGVLTPSS KRFQPFQQFG RDVSDFTDSV RDPKTSEILD ISPCSFGGVS
                    __________________________________SD1_____________________ ___SD2____
```

```
YP_009724390  597  VITPGTNTSN QVAVLYQDVN CTEVPVAIHA DQLTPTWRVY STGSNVFQTR AGCLIGAEHV
QHS34546.1    596  VITPGTNTSN QVAVLYQDVN CTEVPVAIHA DQLTPTWRVY STGSNVFQTR AGCLIGAEHV
QIS60906.1    597  VITPGTNTSN QVAVLYQDVN CTEVPVAIHA DQLTPTWRVY STGSNVFQTR AGCLIGAEHV
QIS60930.1    597  VITPGTNTSN QVAVLYQDVN CTEVPVAIHA DQLTPTWRVY STGSNVFQTR AGCLIGAEHV
QIS61338.1    597  VITPGTNTSN QVAVLYQDVN CTEVPVAIHA DQLTPTWRVY STGSNVFQTR AGCLIGAEHV
QIS60489.1    597  VITPGTNTSN QVAVLYQDVN CTEVPVAIHA DQLTPTWRVY STGSNVFQTR AGCLIGAEHV
QIS30625.1    597  VITPGTNTSN QVAVLYQDVN CTEVPVAIHA DQLTPTWRVY STGSNVFQTR AGCLIGAEHV
QIS30335.1    597  VITPGTNTSN QVAVLYQDVN CTEVPVAIHA DQLTPTWRVY STGSNVFQTR AGCLIGAEHV
QIC53204.1    597  VITPGTNTSN QVAVLYQDVN CTEVPVAIHA DQLTPTWRVY STGSNVFQTR AGCLIGAEHV
QHW06059.1    597  VITPGTNTSN QVAVLYQDVN CTEVPVAIHA DQLTPTWRVY STGSNVFQTR AGCLIGAEHV
QHZ00379.1    597  VITPGTNTSN QVAVLYQDVN CTEVPVAIHA DQLTPTWRVY STGSNVFQTR AGCLIGAEHV
QHR84449.1    597  VITPGTNTSN QVAVLYQDVN CTEVPVAIHA DQLTPTWRVY STGSNVFQTR AGCLIGAEHV
QIJ96493.1    597  VITPGTNTSN QVAVLYQDVN CTEVPVAIHA DQLTPTWRVY STGSNVFQTR AGCLIGAEHV
QIA98583.1    597  VITPGTNTSN QVAVLYQDVN CTEVPVAIHA DQLTPTWRVY STGSNVFQTR AGCLIGAEHV
QII57278.1    597  VITPGTNTSN QVAVLYQDVN CTEVPVAIHA DQLTPTWRVY STGSNVFQTR AGCLIGAEHV
QIA20044.1    597  VITPGTNTSN QVAVLYQDVN CTEVPVAIHA DQLTPTWRVY STGSNVFQTR AGCLIGAEHV
QIS60546.1    597  VITPGTNTSN QVAVLYQDVN CTEVPVAIHA DQLTPTWRVY STGSNVFQTR AGCLIGAEHV
QIS60582.1    597  VITPGTNTSN QVAVLYQDVN CTEVPVAIHA DQLTPTWRVY STGSNVFQTR AGCLIGAEHV
QIS61254.1    597  VITPGTNTSN QVAVLYQDVN CTEVPVAIHA DQLTPTWRVY STGSNVFQTR AGCLIGAEHV
QIS61422.1    597  VITPGTNTSN QVAVLYQDVN CTEVPVAIHA DQLTPTWRVY STGSNVFQTR AGCLIGAEHV
QIS60978.1    597  VITPGTNTSN QVAVLYQDVN CTEVPVAIHA DQLTPTWRVY STGSNVFQTR AGCLIGAEHV
QIO04367.1    597  VITPGTNTSN QVAVLYQDVN CTEVPVAIHA DQLTPTWRVY STGSNVFQTR AGCLIGAEHV
QIS30295.1    597  VITPGTNTSN QVAVLYQGVN CTEVPVAIHA DQLTPTWRVY STGSNVFQTR AGCLIGAEHV
QIS30615.1    597  VITPGTNTSN QVAVLYQGVN CTEVPVAIHA DQLTPTWRVY STGSNVFQTR AGCLIGAEHV
QIK50427.1    597  VITPGTNTSN QVAVLYQGVN CTEVPVAIHA DQLTPTWRVY STGSNVFQTR AGCLIGAEHV
QIS30165.1    597  VITPGTNTSN QVAVLYQDVN CTEVPVAIHA DQLTPTWRVY STGSNVFQTR AGCLIGAEHV
SARS-CoV      583  VITPGTNASS EVAVLYQDVN CTDVSTAIHA DQLTPAWRIY STGNNVFQTQ AGCLIGAEHV
                   ___________________________________________________________
                                             SD2
```

```
YP_009724390  657  NNSYECDIPI GAGICASYQT QTNSPRRARS VASQSIIAYT MSLGAENSVA YSNNSIAIPT
QHS34546.1    656  NNSYECDIPI GAGICASYQT QTNSPRRARS VASQSIIAYT MSLGAENSVA YSNNSIAIPT
QIS60906.1    657  NNSYECDIPI GAGICASYQT QTNSPRRARS VASQSIIAYT MSLGAENSVA YSNNSIAIPT
QIS60930.1    657  NNSYECDIPI GAGICASYQT QTNSPRRARS VASQSIIAYT MSLGAENSVA YSNNSIAIPT
QIS61338.1    657  NNSYECDIPI GAGICASYQT QTNSPRRARS VASQSIIAYT MSLGAENSVA YSNNSIAIPT
QIS60489.1    657  NNSYECDIPI GAGICASYQT QTNSPRRARS VASQSIIAYT MSLGAENSVA YSNNSIAIPT
QIS30625.1    657  NNSYECDIPI GAGICASYQT QTNSPRRARS VASQSIIAYT MSLGAENSVA YSNNSIAIPT
QIS30335.1    657  NNSYECDIPI GAGICASYQT QTNSPRRARS VASQSIIAYT MSLGAENSVA YSNNSIAIPT
QIC53204.1    657  NNSYECDIPI GAGICASYQT QTNSPRRARS VASQSIIAYT MSLGAENSVA YSNNSIAIPT
QHW06059.1    657  NNSYECDIPI GAGICASYQT QTNSPRRARS VASQSIIAYT MSLGAENSVA YSNNSIAIPT
QHZ00379.1    657  NNSYECDIPI GAGICASYQT QTNSPRRARS VASQSIIAYT MSLGAENSVA YSNNSIAIPT
QHR84449.1    657  NNSYECDIPI GAGICASYQT QTNSPRRARS VASQSIIAYT MSLGAENSVA YSNNSIAIPT
QIJ96493.1    657  NNSYECDIPI GAGICASYQT QTNSPRRARS VASQSIIAYT MSLGAENSVA YSNNSIAIPT
QIA98583.1    657  NNSYECDIPI GAGICASYQT QTNSPRRARS VASQSIIAYT MSLGAENSVA YSNNSIAIPT
QII57278.1    657  NNSYECDIPI GAGICASYQT QTNSPRRARS VASQSIIAYT MSLGAENSVA YSNNSIAIPT
QIA20044.1    657  NNSYECDIPI GAGICASYQT QTNSPRRARS VASQSIIAYT MSLGAENSVA YSNNSIAIPT
QIS60546.1    657  NNSYECDIPI GAGICASYQT QTNSPRRARS VASQSIIAYT MSLGAENSVA YSNNSIAIPT
QIS60582.1    657  NNSYECDIPI GAGICASYQT QTNSPRRARS VASQSIIAYT MSLGAENSVA YSNNSIAIPT
QIS61254.1    657  NNSYECDIPI GAGICASYQT QTNSPRRARS VASQSIIAYT MSLGAENSVA YSNNSIAIPT
QIS61422.1    657  NNSYECDIPI GAGICASYQT QTNSPRRARS VASQSIIAYT MSLGAENSVA YSNNSIAIPT
QIS60978.1    657  NNSYECDIPI GAGICASYQT QTNSPRRARS VASQSIIAYT MSLGAENSVA YSNNSIAIPT
QIO04367.1    657  NNSYECDIPI GAGICASYQT QTNSPRRARS VASQSIIAYT MSLGAENSVA YSNNSIAIPT
QIS30295.1    657  NNSYECDIPI GAGICASYQT QTNSPRRARS VASQSIIAYT MSLGAENSVA YSNNSIAIPT
QIS30615.1    657  NNSYECDIPI GAGICASYQT QTNSPRRARS VASQSIIAYT MSLGAENSVA YSNNSIAIPT
QIK50427.1    657  NNSYECDIPI GAGICASYQT QTNSPRRARS VASQSIIAYT MSLGAENSVA YSNNSIAIPT
QIS30165.1    657  NNSYECDIPI GAGICASYQT QTNSPRRARS VASQSIIAYT MSLGAENSVA YSNNSIAIPT
SARS-CoV      643  DTSYECDIPI GAGICASYHT VS----LLRS TSQKSIVAYT MSLGADSSIA YSNNTIAIPT
                   _____________________ __________
                           SD2              S1/S2
```

| | | | | | | | |
|---|---|---|---|---|---|---|---|
| YP_009724390 | 717 | NFTISVTTEI | LPVSMTKTSV | DCTMYICGDS | TECSNLLLQY | GSFCTQLNRA | LTGIAVEQDK |
| QHS34546.1 | 716 | NFTISVTTEI | LPVSMTKTSV | DCTMYICGDS | TECSNLLLQY | GSFCTQLNRA | LTGIAVEQDK |
| QIS60906.1 | 717 | NFTISVTTEI | LPVSMTKTSV | DCTMYICGDS | TECSNLLLQY | GSFCTQLNRA | LTGIAVEQDK |
| QIS60930.1 | 717 | NFTISVTTEI | LPVSMTKTSV | DCTMYICGDS | TECSNLLLQY | GSFCTQLNRA | LTGIAVEQDK |
| QIS61338.1 | 717 | NFTISVTTEI | LPVSMTKTSV | DCTMYICGDS | TECSNLLLQY | GSFCTQLNRA | LTGIAVEQDK |
| QIS60489.1 | 717 | NFTISVTTEI | LPVSMTKTSV | DCTMYICGDS | TECSNLLLQY | GSFCTQLNRA | LTGIAVEQDK |
| QIS30625.1 | 717 | NFTISVTTEI | LPVSMTKTSV | DCTMYICGDS | TECSNLLLQY | GSFCTQLNRA | LTGIAVEQDK |
| QIS30335.1 | 717 | NFTISVTTEI | LPVSMTKTSV | DCTMYICGDS | TECSNLLLQY | GSFCTQLNRA | LTGIAVEQDK |
| QIC53204.1 | 717 | NFTISVTTEI | LPVSMTKTSV | DCTMYICGDS | TECSNLLLQY | GSFCTQLNRA | LTGIAVEQDK |
| QHW06059.1 | 717 | NFTISVTTEI | LPVSMTKTSV | DCTMYICGDS | TECSNLLLQY | GSFCTQLNRA | LTGIAVEQDK |
| QHZ00379.1 | 717 | NFTISVTTEI | LPVSMTKTSV | DCTMYICGDS | TECSNLLLQY | GSFCTQLNRA | LTGIAVEQDK |
| QHR84449.1 | 717 | NFTISVTTEI | LPVSMTKTSV | DCTMYICGDS | TECSNLLLQY | GSFCTQLNRA | LTGIAVEQDK |
| QIJ96493.1 | 717 | NFTISVTTEI | LPVSMTKTSV | DCTMYICGDS | TECSNLLLQY | GSFCTQLNRA | LTGIAVEQDK |
| QIA98583.1 | 717 | NFTISVTTEI | LPVSMTKTSV | DCTMYICGDS | TECSNLLLQY | GSFCTQLNRA | LTGIAVEQDK |
| QII57278.1 | 717 | NFTISVTTEI | LPVSMTKTSV | DCTMYICGDS | TECSNLLLQY | GSFCTQLNRA | LTGIAVEQDK |
| QIA20044.1 | 717 | NFTISVTTEI | LPVSMTKTSV | DCTMYICGDS | TECSNLLLQY | GSFCTQLNRA | LTGIAVEQDK |
| QIS60546.1 | 717 | NFTISVTTEI | LPVSMTKTSV | DCTMYICGDS | TECSNLLLQY | GSFCTQLNRA | LTGIAVEQDK |
| QIS60582.1 | 717 | NFTISVTTEI | LPVSMTKTSV | DCTMYICGDS | TECSNLLLQY | GSFCTQLNRA | LTGIAVEQDK |
| QIS61254.1 | 717 | NFTISVTTEI | LPVSMTKTSV | DCTMYICGDS | TECSNLLLQY | GSFCTQLNRA | LTGIAVEQDK |
| QIS61422.1 | 717 | NFTISVTTEI | LPVSMTKTSV | DCTMYICGDS | TECSNLLLQY | GSFCTQLNRA | LTGIAVEQDK |
| QIS60978.1 | 717 | NFTISVTTEI | LPVSMTKTSV | DCTMYICGDS | TECSNLLLQY | GSFCTQLNRA | LTGIAVEQDK |
| QIO04367.1 | 717 | NFTISVTTEI | LPVSMTKTSV | DCTMYICGDS | TECSNLLLQY | GSFCTQLNRA | LTGIAVEQDK |
| QIS30295.1 | 717 | NFTISVTTEI | LPVSMTKTSV | DCTMYICGDS | TECSNLLLQY | GSFCTQLNRA | LTGIAVEQDK |
| QIS30615.1 | 717 | NFTISVTTEI | LPVSMTKTSV | DCTMYICGDS | TECSNLLLQY | GSFCTQLNRA | LTGIAVEQDK |
| QIK50427.1 | 717 | NFTISVTTEI | LPVSMTKTSV | DCTMYICGDS | TECSNLLLQY | GSFCTQLNRA | LTGIAVEQDK |
| QIS30165.1 | 717 | NFTISVTTEI | LPVSMTKTSV | DCTMYICGDS | TECSNLLLQY | GSFCTQLNRA | LTGIAVEQDK |
| SARS-CoV | 699 | NFSISITTEV | MPVSMAKTSV | DCNMYICGDS | TECANLLLQY | GSFCTQLNRA | LSGIAAEQDR |

| | | | | | | | |
|---|---|---|---|---|---|---|---|
| YP_009724390 | 777 | NTQEVFAQVK | QIYKTPPIKD | FGGFNFSQIL | PDPSKPSKRS | FIEDLLFNKV | TLADAGFIKQ |
| QHS34546.1 | 776 | NTQEVFAQVK | QIYKTPPIKD | FGGFNFSQIL | PDPSKPSKRS | FIEDLLFNKV | TLADAGFIKQ |
| QIS60906.1 | 777 | NTQEVFAQVK | QIYKTPPIKD | FGGFNFSQIL | PDPSKPSKRS | FIEDLLFNKV | TLADAGFIKQ |
| QIS60930.1 | 777 | NTQEVFAQVK | QIYKTPPIKD | FGGFNFSQIL | PDPSKPSKRS | FIEDLLFNKV | TLADAGFIKQ |
| QIS61338.1 | 777 | NTQEVFAQVK | QIYKTPPIKD | FGGFNFSQIL | PDPSKPSKRS | FIEDLLFNKV | TLADAGFIKQ |
| QIS60489.1 | 777 | NTQEVFAQVK | QIYKTPPIKD | FGGFNFSQIL | PDPSKPSKRS | FIEDLLFNKV | TLADAGFIKQ |
| QIS30625.1 | 777 | NTQEVFAQVK | QIYKTPPIKD | FGGFNFSQIL | PDPSKPSKRS | FIEDLLFNKV | TLADAGFIKQ |
| QIS30335.1 | 777 | NTQEVFAQVK | QIYKTPPIKD | FGGFNFSQIL | PDPSKPSKRS | FIEDLLFNKV | TLADAGFIKQ |
| QIC53204.1 | 777 | NTQEVFAQVK | QIYKTPPIKD | CGGFNFSQIL | PDPSKPSKRS | FIEDLLFNKV | TLADAGFIKQ |
| QHW06059.1 | 777 | NTQEVFAQVK | QIYKTPPIKD | FGGFNFSQIL | PDPSKPSKRS | FIEDLLFNKV | TLADAGFIKQ |
| QHZ00379.1 | 777 | NTQEVFAQVK | QIYKTPPIKD | FGGFNFSQIL | PDPSKPSKRS | FIEDLLFNKV | TLADAGFIKQ |
| QHR84449.1 | 777 | NTQEVFAQVK | QIYKTPPIKD | FGGFNFSQIL | PDPSKPSKRS | FIEDLLFNKV | TLADAGFIKQ |
| QIJ96493.1 | 777 | NTQEVFAQVK | QIYKTPPIKD | FGGFNFSQIL | PDPSKPSKRS | FIEDLLFNKV | TLADAGFIKQ |
| QIA98583.1 | 777 | NTQEVFAQVK | QIYKTPPIKD | FGGFNFSQIL | PDPSKPSKRS | FIEDLLFNKV | TLADAGFIKQ |
| QII57278.1 | 777 | NTQEVFAQVK | QIYKTPPIKD | FGGFNFSQIL | PDPSKPSKRS | FIEDLLFNKV | TLADAGFIKQ |
| QIA20044.1 | 777 | NTQEVFAQVK | QIYKTPPIKD | FGGFNFSQIL | PDPSKPSKRS | FIEDLLFNKV | TLADAGFIKQ |
| QIS60546.1 | 777 | NTQEVFAQVK | QIYKTPPIKD | FGGFNFSQIL | PDPSKPSKRS | FIEDLLFNKV | TLADAGFIKQ |
| QIS60582.1 | 777 | NTQEVFAQVK | QIYKTPPIKD | FGGFNFSQIL | PDPSKPSKRS | FIEDLLFNKV | TLADAGFIKQ |
| QIS61254.1 | 777 | NTQEVFAQVK | QIYKTPPIKD | FGGFNFSQIL | PDPSKPSKRS | FIEDLLFNKV | TLADAGFIKQ |
| QIS61422.1 | 777 | NTQEVFAQVK | QIYKTPPIKD | FGGFNFSQIL | PDPSKPSKRS | FIEDLLFNKV | TLADAGFIKQ |
| QIS60978.1 | 777 | NTQEVFAQVK | QIYKTPPIKD | FGGFNFSQIL | PDPSKPSKRS | FIEDLLFNKV | TLADAGFIKQ |
| QIO04367.1 | 777 | NTQEVFAQVK | QIYKTPPIKD | FGGFNFSQIL | PDPSKPSKRS | FIEDLLFNKV | TLADAGFIKQ |
| QIS30295.1 | 777 | NTQEVFAQVK | QIYKTPPIKD | FGGFNFSQIL | PDPSKPSKRS | FIEDLLFNKV | TLADAGFIKQ |
| QIS30615.1 | 777 | NTQEVFAQVK | QIYKTPPIKD | FGGFNFSQIL | PDPSKPSKRS | FIEDLLFNKV | TLADAGFIKQ |
| QIK50427.1 | 777 | NTQEVFAQVK | QIYKTPPIKD | FGGFNFSQIL | PDPSKPSKRS | FIEDLLFNKV | TLADAGFIKQ |
| QIS30165.1 | 777 | NTQEVFAQVK | QIYKTPPIKD | FGGFNFSQIL | PDPSKPSKRS | FIEDLLFNKV | TLADAGFIKQ |
| SARS-CoV | 759 | NTREVFAQVK | QMYKTPTLKY | FGGFNFSQIL | PDPLKPTKRS | FIEDLLFNKV | TLADAGFMKQ |

S2′ FP

| | | | | | | | |
|---|---|---|---|---|---|---|---|
| YP_009724390 | 837 | YGDCLGDIAA | RDLICAQKFN | GLTVLPPLLT | DEMIAQYTSA | LLAGTITSGW | TFGAGAALQI |
| QHS34546.1 | 836 | YGDCLGDIAA | RDLICAQKFN | GLTVLPPLLT | DEMIAQYTSA | LLAGTITSGW | TFGAGAALQI |
| QIS60906.1 | 837 | YGDCLGDIAA | RDLICAQKFN | GLTVLPPLLT | DEMIAQYTSA | LLAGTITSGW | TFGAGAALQI |
| QIS60930.1 | 837 | YGDCLGDIAA | RDLICAQKFN | GLTVLPPLLT | DEMIAQYTSA | LLAGTITSGW | TFGAGAALQI |

| | | | | | | | |
|---|---|---|---|---|---|---|---|
| QIS61338.1 | 837 | YGDCLGDIAA | RDLICAQKFN | GLTVLPPLLT | DEMIAQYTSA | LLAGTITSGW | TFGAGAALQI |
| QIS60489.1 | 837 | YGDCLGDIAA | RDLICAQKFN | GLTVLPPLLT | DEMIAQYTSA | LLAGTITSGW | TFGAGAALQI |
| QIS30625.1 | 837 | YGDCLGDIAA | RDLICAQKFN | GLTVLPPLLT | DEMIAQYTSA | LLAGTITSGW | TFGAGAALQI |
| QIS30335.1 | 837 | YGDCLGDIAA | RDLICAQKFN | GLTVLPPLLT | DEMIAQYTSA | LLAGTITSGW | TFGAGAALQI |
| QIC53204.1 | 837 | YGDCLGDIAA | RDLICAQKFN | GLTVLPPLLT | DEMIAQYTSA | LLAGTITSGW | TFGAGAALQI |
| QHW06059.1 | 837 | YGDCLGDIAA | RDLICAQKFN | GLTVLPPLLT | DEMIAQYTSA | LLAGTITSGW | TFGAGAALQI |
| QHZ00379.1 | 837 | YGDCLGDIAA | RDLICAQKFN | GLTVLPPLLT | DEMIAQYTSA | LLAGTITSGW | TFGAGAALQI |
| QHR84449.1 | 837 | YGDCLGDIAA | RDLICAQKFN | GLTVLPPLLT | DEMIAQYTSA | LLAGTITSGW | TFGAGAALQI |
| QIJ96493.1 | 837 | YGDCLGDIAA | RDLICAQKFN | GLTVLPPLLT | DEMIAQYTSA | LLAGTITSGW | TFGAGAALQI |
| QIA98583.1 | 837 | YGDCLGDIAA | RDLICAQKFN | GLTVLPPLLT | DEMIAQYTSA | LLAGTITSGW | TFGAGAALQI |
| QII57278.1 | 837 | YGDCLGDIAA | RDLICAQKFN | GLTVLPPLLT | DEMIAQYTSA | LLAGTITSGW | TFGAGAALQI |
| QIA20044.1 | 837 | YGDCLGDIAA | RDLICAQKFN | GLTVLPPLLT | DEMIAQYTSA | LLAGTITSGW | TFGAGAALQI |
| QIS60546.1 | 837 | YGDCLGDIAA | RDLICAQKFN | GLTVLPPLLT | DEMIAQYTSA | LLAGTITSGW | TFGAGAALQI |
| QIS60582.1 | 837 | YGDCLGDIAA | RDLICAQKFN | GLTVLPPLLT | DEMIAQYTSA | LLAGTITSGW | TFGAGAALQI |
| QIS61254.1 | 837 | YGDCLGDIAA | RDLICAQKFN | GLTVLPPLLT | DEMIAQYTSA | LLAGTITSGW | TFGAGAALQI |
| QIS61422.1 | 837 | YGDCLGDIAA | RDLICAQKFN | GLTVLPPLLT | DEMIAQYTSA | LLAGTITSGW | TFGAGAALQI |
| QIS60978.1 | 837 | YGDCLGDIAA | RDLICAQKFN | GLTVLPPLLT | DEMIAQYTSA | LLAGTITSGW | TFGAGAALQI |
| QIO04367.1 | 837 | YGDCLGDIAA | RDLICAQKFN | GLTVLPPLLT | DEMIAQYTSA | LLAGTITSGW | TFGAGAALQI |
| QIS30295.1 | 837 | YGDCLGDIAA | RDLICAQKFN | GLTVLPPLLT | DEMIAQYTSA | LLAGTITSGW | TFGAGAALQI |
| QIS30615.1 | 837 | YGDCLGDIAA | RDLICAQKFN | GLTVLPPLLT | DEMIAQYTSA | LLAGTITSGW | TFGAGAALQI |
| QIK50427.1 | 837 | YGDCLGDIAA | RDLICAQKFN | GLTVLPPLLT | DEMIAQYTSA | LLAGTITSGW | TFGAGAALQI |
| QIS30165.1 | 837 | YGDCLGDIAA | RDLICAQKFN | GLTVLPPLLT | DEMIAQYTSA | LLAGTITSGW | TFGAGAALQI |
| SARS-CoV | 819 | YGECLGDINA | RDLICAQKFN | GLTVLPPLLT | DDMIAAYTAA | LVSGTATAGW | TFGAGAALQI |

CR

| | | | | | | | |
|---|---|---|---|---|---|---|---|
| YP_009724390 | 897 | PFAMQMAYRF | NGIGVTQNVL | YENQKLIANQ | FNSAIGKIQD | SLSSTASALG | KLQDVVNQNA |
| QHS34546.1 | 896 | PFAMQMAYRF | NGIGVTQNVL | YENQKLIANQ | FNSAIGKIQD | SLSSTASALG | KLQDVVNQNA |
| QIS60906.1 | 897 | PFAMQMAYRF | NGIGVTQNVL | YENQKLIANQ | FNSAIGKIQD | SLSSTASALG | KLQDVVNQNA |
| QIS60930.1 | 897 | PFAMQMAYRF | NGIGVTQNVL | YENQKLIANQ | FNSAIGKIQD | SLSSTASALG | KLQDVVNQNA |
| QIS61338.1 | 897 | PFAMQMAYRF | NGIGVTQNVL | YENQKLIANQ | FNSAIGKIQD | SLSSTASALG | KLQDVVNQNA |
| QIS60489.1 | 897 | PFAMQMAYRF | NGIGVTQNVL | YENQKLIANQ | FNSAIGKIQD | SLSSTASALG | KLQDVVNQNA |
| QIS30625.1 | 897 | PFAMQMAYRF | NGIGVTQNVL | YENQKLIANQ | FNSAIGKIQD | SLSSTASALG | KLQDVVNQNA |
| QIS30335.1 | 897 | PFAMQMAYRF | NGIGVTQNVL | YENQKLIANQ | FNSAIGKIQD | SLSSTASALG | KLQDVVNQNA |
| QIC53204.1 | 897 | PFAMQMAYRF | NGIGVTQNVL | YENQKLIANQ | FNSAIGKIQD | SLSSTASALG | KLQDVVNQNA |
| QHW06059.1 | 897 | PFAMQMAYRF | NGIGVTQNVL | YENQKLIANQ | FNSAIGKIQD | SLSSTASALG | KLQDVVNQNA |
| QHZ00379.1 | 897 | PFAMQMAYRF | NGIGVTQNVL | YENQKLIANQ | FNSAIGKIQD | SLSSTASALG | KLQDVVNQNA |
| QHR84449.1 | 897 | PFAMQMAYRF | NGIGVTQNVL | YENQKLIANQ | FNSAIGKIQD | SLSSTASALG | KLQDVVNQNA |
| QIJ96493.1 | 897 | PFAMQMAYRF | NGIGVTQNVL | YENQKLIANQ | FNSAIGKIQD | SLSSTASALG | KLQDVVNQNA |
| QIA98583.1 | 897 | PFAMQMAYRF | NGIGVTQNVL | YENQKLIANQ | FNSVIGKIQD | SLSSTASALG | KLQDVVNQNA |
| QII57278.1 | 897 | PFAMQMAYRF | NGIGVTQNVL | YENQKLIANQ | FNSAIGKIQD | SLSSTASALG | KLQDVVNQNA |
| QIA20044.1 | 897 | PFAMQMAYRF | NGIGVTQNVL | YENQKLIANQ | FNSAIGKIQD | SLSSTASALG | KLQDVVNQNA |
| QIS60546.1 | 897 | PFAMQMAYRF | NGIGVTQNVL | YENQKLIANQ | FNSAIGKIQD | SLSSTASALG | KLQDVVNQNA |
| QIS60582.1 | 897 | PFAMQMAYRF | NGIGVTQNVL | YENQKLIANQ | FNSAIGKIQD | SLSSTASALG | KLQDVVNQNA |
| QIS61254.1 | 897 | PFAMQMAYRF | NGIGVTQNVL | YENQKLIANQ | FNSAIGKIQD | SLSSTASALG | KLQDVVNQNA |
| QIS61422.1 | 897 | PFAMQMAYRF | NGIGVTQNVL | YENQKLIANQ | FNSAIGKIQD | SLSSTASALG | KLQDVVNQNA |
| QIS60978.1 | 897 | PFAMQMAYRF | NGIGVTQNVL | YENQKLIANQ | FNSAIGKIQD | SLSSTASALG | KLQDVVNQNA |
| QIO04367.1 | 897 | PFAMQMAYRF | NGIGVTQNVL | YENQKLIANQ | FNSAIGKIQD | SLSSTASALG | KLQDVVNQNA |
| QIS30295.1 | 897 | PFAMQMAYRF | NGIGVTQNVL | YENQKLIANQ | FNSAIGKIQD | SLSSTASALG | KLQDVVNQNA |
| QIS30615.1 | 897 | PFAMQMAYRF | NGIGVTQNVL | YENQKLIANQ | FNSAIGKIQY | SLSSTASALG | KLQDVVNQNA |
| QIK50427.1 | 897 | PFAMQMAYRF | NGIGVTQNVL | YENQKLIANQ | FNSAIGKIQD | SLSSTASALG | KLQDVVNQNA |
| QIS30165.1 | 897 | PFAMQMAYRF | NGIGVTQNVL | YENQKLIANQ | FNSAIGKIQD | SLSSTASALG | KLQDVVNQNA |
| SARS-CoV | 879 | PFAMQMAYRF | NGIGVTQNVL | YENQKQIANQ | FNKAISQIQE | SLTTTSTALG | KLQDVVNQNA |

CR HR1

| | | | | | | | |
|---|---|---|---|---|---|---|---|
| YP_009724390 | 957 | QALNTLVKQL | SSNFGAISSV | LNDILSRLDK | VEAEVQIDRL | ITGRLQSLQT | YVTQQLIRAA |
| QHS34546.1 | 956 | QALNTLVKQL | SSNFGAISSV | LNDILSRLDK | VEAEVQIDRL | ITGRLQSLQT | YVTQQLIRAA |
| QIS60906.1 | 957 | QALNTLVKQL | SSNFGAISSV | LNDILSRLDK | VEAEVQIDRL | ITGRLQSLQT | YVTQQLIRAA |
| QIS60930.1 | 957 | QALNTLVKQL | SSNFGAISSV | LNDILSRLDK | VEAEVQIDRL | ITGRLQSLQT | YVTQQLIRAA |
| QIS61338.1 | 957 | QALNTLVKQL | SSNFGAISSV | LNDILSRLDK | VEAEVQIDRL | ITGRLQSLQT | YVTQQLIRAA |
| QIS60489.1 | 957 | QALNTLVKQL | SSNFGAISSV | LNDILSRLDK | VEAEVQIDRL | ITGRLQSLQT | YVTQQLIRAA |

| | | | | | | | |
|---|---|---|---|---|---|---|---|
| QIS30625.1 | 957 | QALNTLVKQL | SSNFGAISSV | LNDILSRLDK | VEAEVQIDRL | ITGRLQSLQT | YVTQQLIRAA |
| QIS30335.1 | 957 | QALNTLVKQL | SSNFGAISSV | LNDILSRLDK | VEAEVQIDRL | ITGRLQSLQT | YVTQQLIRAA |
| QIC53204.1 | 957 | QALNTLVKQL | SSNFGAISSV | LNDILSRLDK | VEAEVQIDRL | ITGRLQSLQT | YVTQQLIRAA |
| QHW06059.1 | 957 | QALNTLVKQL | SSNFGAISSV | LNDILSRLDK | VEAEVQIDRL | ITGRLQSLQT | YVTQQLIRAA |
| QHZ00379.1 | 957 | QALNTLVKQL | SSNFGAISSV | LNDILSRLDK | VEAEVQIDRL | ITGRLQSLQT | YVTQQLIRAA |
| QHR84449.1 | 957 | QALNTLVKQL | SSNFGAISSV | LNDILSRLDK | VEAEVQIDRL | ITGRLQSLQT | YVTQQLIRAA |
| QIJ96493.1 | 957 | QALNTLVKQL | SSNFGAISSV | LNDILSRLDK | VEAEVQIDRL | ITGRLQSLQT | YVTQQLIRAA |
| QIA98583.1 | 957 | QALNTLVKQL | SSNFGAISSV | LNDILSRLDK | VEAEVQIDRL | ITGRLQSLQT | YVTQQLIRAA |
| QII57278.1 | 957 | QALNTLVKQL | SSNFGAISSV | LNDILSRLDK | VEAEVQIDRL | ITGRLQSLQT | YVTQQLIRAA |
| QIA20044.1 | 957 | QALNTLVKQL | SSNFGAISSV | LNDILSRLDK | VEAEVQIDRL | ITGRLQSLQT | YVTQQLIRAA |
| QIS60546.1 | 957 | QALNTLVKQL | SSNFGAISSV | LNDILSRLDK | VEAEVQIDRL | ITGRLQSLQT | YVTQQLIRAA |
| QIS60582.1 | 957 | QALNTLVKQL | SSNFGAISSV | LNDILSRLDK | VEAEVQIDRL | ITGRLQSLQT | YVTQQLIRAA |
| QIS61254.1 | 957 | QALNTLVKQL | SSNFGAISSV | LNDILSRLDK | VEAEVQIDRL | ITGRLQSLQT | YVTQQLIRAA |
| QIS61422.1 | 957 | QALNTLVKQL | SSNFGAISSV | LNDILSRLDK | VEAEVQIDRL | ITGRLQSLQT | YVTQQLIRAA |
| QIS60978.1 | 957 | QALNTLVKQL | SSNFGAISSV | LNDILSRLDK | VEAEVQIDRL | ITGRLQSLQT | YVTQQLIRAA |
| QIO04367.1 | 957 | QALNTLVKQL | SSNFGAISSV | LNDILSRLDK | VEAEVQIDRL | ITGRLQSLQT | YVTQQLIRAA |
| QIS30295.1 | 957 | QALNTLVKQL | SSNFGAISSV | LNDILSRLDK | VEAEVQIDRL | ITGRLQSLQT | YVTQQLIRAA |
| QIS30615.1 | 957 | QALNTLVKQL | SSNFGAISSV | LNDILSRLDK | VEAEVQIDRL | ITGRLQSLQT | YVTQQLIRAA |
| QIK50427.1 | 957 | QALNTLVKQL | SSNFGAISSV | LNDILSRLDK | VEAEVQIDRL | ITGRLQSLQT | YVTQQLIRAA |
| QIS30165.1 | 957 | QALNTLVKQL | SSNFGAISSV | LNDILSRLDK | VEAEVQIDRL | ITGRLQSLQT | YVTQQLIRAA |
| SARS-CoV | 939 | QALNTLVKQL | SSNFGAISSV | LNDILSRLDK | VEAEVQIDRL | ITGRLQSLQT | YVTQQLIRAA |

HR1 CH

| | | | | | | | |
|---|---|---|---|---|---|---|---|
| YP_009724390 | 1017 | EIRASANLAA | TKMSECVLGQ | SKRVDFCGKG | YHLMSFPQSA | PHGVVFLHVT | YVPAQEKNFT |
| QHS34546.1 | 1016 | EIRASANLAA | TKMSECVLGQ | SKRVDFCGKG | YHLMSFPQSA | PHGVVFLHVT | YVPAQEKNFT |
| QIS60906.1 | 1017 | EIRASANLAA | TKMSECVLGQ | SKRVDFCGKG | YHLMSFPQSA | PHGVVFLHVT | YVPAQEKNFT |
| QIS60930.1 | 1017 | EIRASANLAA | TKMSECVLGQ | SKRVDFCGKG | YHLMSFPQSA | PHGVVFLHVT | YVPAQEKNFT |
| QIS61338.1 | 1017 | EIRASANLAA | TKMSECVLGQ | SKRVDFCGKG | YHLMSFPQSA | PHGVVFLHVT | YVPAQEKNFT |
| QIS60489.1 | 1017 | EIRASANLAA | TKMSECVLGQ | SKRVDFCGKG | YHLMSFPQSA | PHGVVFLHVT | YVPAQEKNFT |
| QIS30625.1 | 1017 | EIRASANLAA | TKMSECVLGQ | SKRVDFCGKG | YHLMSFPQSA | PHGVVFLHVT | YVPAQEKNFT |
| QIS30335.1 | 1017 | EIRASANLAA | TKMSECVLGQ | SKRVDFCGKG | YHLMSFPQSA | PHGVVFLHVT | YVPAQEKNFT |
| QIC53204.1 | 1017 | EIRASANLAA | TKMSECVLGQ | SKRVDFCGKG | YHLMSFPQSA | PHGVVFLHVT | YVPAQEKNFT |
| QHW06059.1 | 1017 | EIRASANLAA | TKMSECVLGQ | SKRVDFCGKG | YHLMSFPQSA | PHGVVFLHVT | YVPAQEKNFT |
| QHZ00379.1 | 1017 | EIRASANLAA | TKMSECVLGQ | SKRVDFCGKG | YHLMSFPQSA | PHGVVFLHVT | YVPAQEKNFT |
| QHR84449.1 | 1017 | EIRASANLAA | TKMSECVLGQ | SKRVDFCGKG | YHLMSFPQSA | PHGVVFLHVT | YVPAQEKNFT |
| QIJ96493.1 | 1017 | EIRASANLAA | TKMSECVLGQ | SKRVDFCGKG | YHLMSFPQSA | PHGVVFLHVT | YVPAQEKNFT |
| QIA98583.1 | 1017 | EIRASANLAA | TKMSECVLGQ | SKRVDFCGKG | YHLMSFPQSA | PHGVVFLHVT | YVPAQEKNFT |
| QII57278.1 | 1017 | EIRASANLAA | TKMSECVLGQ | SKRVDFCGKG | YHLMSFPQSA | PHGVVFLHVT | YVPAQEKNFT |
| QIA20044.1 | 1017 | EIRASANLAA | TKMSECVLGQ | SKRVDFCGKG | YHLMSFPQSA | PHGVVFLHVT | YVPAQEKNFT |
| QIS60546.1 | 1017 | EIRASANLAA | TKMSECVLGQ | SKRVDFCGKG | YHLMSFPQSA | PHGVVFLHVT | YVPAQEKNFT |
| QIS60582.1 | 1017 | EIRASANLAA | TKMSECVLGQ | SKRVDFCGKG | YHLMSFPQSA | PHGVVFLHVT | YVPAQEKNFT |
| QIS61254.1 | 1017 | EIRASANLAA | TKMSECVLGQ | SKRVDFCGKG | YHLMSFPQSA | PHGVVFLHVT | YVPAQEKNFT |
| QIS61422.1 | 1017 | EIRASANLAA | TKMSECVLGQ | SKRVDFCGKG | YHLMSFPQSA | PHGVVFLHVT | YVPAQEKNFT |
| QIS60978.1 | 1017 | EIRASANLAA | TKMSECVLGQ | SKRVDFCGKG | YHLMSFPQSA | PHGVVFLHVT | YVPAQEKNFT |
| QIO04367.1 | 1017 | EIRASANLAA | TKMSECVLGQ | SKRVDFCGKG | YHLMSFPQSA | PHGVVFLHVT | YVPAQEKNFT |
| QIS30295.1 | 1017 | EIRASANLAA | TKMSECVLGQ | SKRVDFCGKG | YHLMSFPQSA | PHGVVFLHVT | YVPAQEKNFT |
| QIS30615.1 | 1017 | EIRASANLAA | TKMSECVLGQ | SKRVDFCGKG | YHLMSFPQSA | PHGVVFLHVT | YVPAQEKNFT |
| QIK50427.1 | 1017 | EIRASANLAA | TKMSECVLGQ | SKRVDFCGKG | YHLMSFPQSA | PHGVVFLHVT | YVPAQEKNFT |
| QIS30165.1 | 1017 | EIRASANLAA | TKMSECVLGQ | SKRVDFCGKG | YHLMSFPQSA | PHGVVFLHVT | YVPAQEKNFT |
| SARS-CoV | 999 | EIRASANLAA | TKMSECVLGQ | SKRVDFCGKG | YHLMSFPQAA | PHGVVFLHVT | YVPSQERNFT |

CH BH SD3

| | | | | | | | |
|---|---|---|---|---|---|---|---|
| YP_009724390 | 1077 | TAPAICHDGK | AHFPREGVFV | SNGTHWFVTQ | RNFYEPQIIT | TDNTFVSGNC | DVVIGIVNNT |
| QHS34546.1 | 1076 | TAPAICHDGK | AHFPREGVFV | SNGTHWFVTQ | RNFYEPQIIT | TDNTFVSGNC | DVVIGIVNNT |
| QIS60906.1 | 1077 | TAPAICHDGK | AHFPREGVFV | SNGTHWFVTQ | RNFYEPQIIT | TDNTFVSGNC | DVVIGIVNNT |
| QIS60930.1 | 1077 | TAPAICHDGK | AHFPREGVFV | SNGTHWFVTQ | RNFYEPQIIT | TDNTFVSGNC | DVVIGIVNNT |
| QIS61338.1 | 1077 | TAPAICHDGK | AHFPREGVFV | SNGTHWFVTQ | RNFYEPQIIT | TDNTFVSGNC | DVVIGIVNNT |
| QIS60489.1 | 1077 | TAPAICHDGK | AHFPREGVFV | SNGTHWFVTQ | RNFYEPQIIT | TDNTFVSGNC | DVVIGIVNNT |
| QIS30625.1 | 1077 | TAPAICHDGK | AHFPREGVFV | SNGTHWFVTQ | RNFYEPQIIT | TDNTFVSGNC | DVVIGIVNNT |

| | | | | | | | |
|---|---|---|---|---|---|---|---|
| QIS30335.1 | 1077 | TAPAICHDGK | AHFPREGVFV | SNGTHWFVTQ | RNFYEPQIIT | TDNTFVSGNC | DVVIGIVNNT |
| QIC53204.1 | 1077 | TAPAICHDGK | AHFPREGVFV | SNGTHWFVTQ | RNFYEPQIIT | TDNTFVSGNC | DVVIGIVNNT |
| QHW06059.1 | 1077 | TAPAICHDGK | AHFPREGVFV | SNGTHWFVTQ | RNFYEPQIIT | TDNTFVSGNC | DVVIGIVNNT |
| QHZ00379.1 | 1077 | TAPAICHDGK | AHFPREGVFV | SNGTHWFVTQ | RNFYEPQIIT | TDNTFVSGNC | DVVIGIVNNT |
| QHR84449.1 | 1077 | TAPAICHDGK | AHFPREGVFV | SNGTHWFVTQ | RNFYEPQIIT | TDNTFVSGNC | DVVIGIVNNT |
| QIJ96493.1 | 1077 | TAPAICHDGK | AHFPREGVFV | SNGTHWFVTQ | RNFYEPQIIT | TDNTFVSGNC | DVVIGIVNNT |
| QIA98583.1 | 1077 | TAPAICHDGK | AHFPREGVFV | SNGTHWFVTQ | RNFYEPQIIT | TDNTFVSGNC | DVVIGIVNNT |
| QII57278.1 | 1077 | TAPAICHDGK | AHFPREGVFV | SNGTHWFVTQ | RNFYEPQIIT | TDNTFVSGNC | DVVIGIVNNT |
| QIA20044.1 | 1077 | TAPAICHDGK | AHFPREGVFV | SNGTHWFVTQ | RNFYEPQIIT | TDNTFVSGNC | DVVIGIVNNT |
| QIS60546.1 | 1077 | TAPAICHDGK | AHFPREGVFV | SNGTHWFVTQ | RNFYEPQIIT | TDNTFVSGNC | DVVIGIVNNT |
| QIS60582.1 | 1077 | TAPAICHDGK | AHFPREGVFV | SNGTHWFVTQ | RNFYEPQIIT | TDNTFVSGNC | DVVIGIVNNT |
| QIS61254.1 | 1077 | TVPAICHDGK | AHFPREGVFV | SNGTHWFVTQ | RNFYEPQIIT | TDNTFVSGNC | DVVIGIVNNT |
| QIS61422.1 | 1077 | TAPAICHDGK | AHFPREGVFV | SNGTHWFVTQ | RNFYEPQIIT | TDNTFVSGNC | DVVIGIVNNT |
| QIS60978.1 | 1077 | TAPAICHDGK | AHFPREGVFV | SNGTHWFVTQ | RNFYEPQIIT | TDNTFVSGNC | DVVIGIVNNT |
| QIO04367.1 | 1077 | TAPAICHDGK | AHFPREGVFV | SNGTHWFVTQ | RNFYEPQIIT | TDNTFVSGNC | DVVIGIVNNT |
| QIS30295.1 | 1077 | TAPAICHDGK | AHFPREGVFV | SNGTHWFVTQ | RNFYEPQIIT | TDNTFVSGNC | DVVIGIVNNT |
| QIS30615.1 | 1077 | TAPAICHDGK | AHFPREGVFV | SNGTHWFVTQ | RNFYEPQIIT | TDNTFVSGNC | DVVIGIVNNT |
| QIK50427.1 | 1077 | TAPAICHDGK | AHFPREGVFV | SNGTHWFVTQ | RNFYEPQIIT | TDNTFVSGNC | DVVIGIVNNT |
| QIS30165.1 | 1077 | TAPAICHDGK | AHFPREGVFV | SNGTHWFVTQ | RNFYEPQIIT | TDNTFVSGNC | DVVIGIVNNT |
| SARS-CoV | 1059 | TAPAICHEGK | AYFPREGVFV | FNGTSWFITQ | RNFFSPQIIT | TDNTFVSGNC | DVVIGIINNT |

SD3

| | | | | | | | |
|---|---|---|---|---|---|---|---|
| YP_009724390 | 1137 | VYDPLQPELD | SFKEELDKYF | KNHTSPDVDL | GDISGINASV | VNIQKEIDRL | NEVAKNLNES |
| QHS34546.1 | 1136 | VYDPLQPELD | SFKEELDKYF | KNHTSPDVDL | GDISGINASV | VNIQKEIDRL | NEVAKNLNES |
| QIS60906.1 | 1137 | VYDPLQPELD | SFKEELDKYF | KNHTSPDVDL | GDISGINASV | VNIQKEIDRL | NEVAKNLNES |
| QIS60930.1 | 1137 | VYDPLQPELD | SFKEELDKYF | KNHTSPDVDL | GDISGINASV | VNIQKEIDRL | NEVAKNLNES |
| QIS61338.1 | 1137 | VYDPLQPELD | SFKEELDKYF | KNHTSPDVDL | GDISGINASV | VNIQKEIDRL | NEVAKNLNES |
| QIS60489.1 | 1137 | VYDPLQPELD | SFKEELDKYF | KNHTSPDVDL | GDISGINASV | VNIQKEIDRL | NEVAKNLNES |
| QIS30625.1 | 1137 | VYDPLQPELD | SFKEELDKYF | KNHTSPDVDL | GDISGINASV | VNIQKEIDRL | NEVAKNLNES |
| QIS30335.1 | 1137 | VYDPLQPELD | SFKEELDKYF | KNHTSPDVDL | GDISGINASV | VNIQKEIDRL | NEVAKNLNES |
| QIC53204.1 | 1137 | VYDPLQPELD | SFKEELDKYF | KNHTSPDVDL | GDISGINASV | VNIQKEIDRL | NEVAKNLNES |
| QHW06059.1 | 1137 | VYDPLQPELD | SFKEELDKYF | KNHTSPDVDL | GDISGINASV | VNIQKEIDRL | NEVAKNLNES |
| QHZ00379.1 | 1137 | VYDPLQPELD | SFKEELDKYF | KNHTSPDVDL | GDISGINASV | VNIQKEIDRL | NEVAKNLNES |
| QHR84449.1 | 1137 | VYDPLQPELD | SFKEELDKYF | KNHTSPDVDL | GDISGINASV | VNIQKEIDRL | NEVAKNLNES |
| QIJ96493.1 | 1137 | VYDPLQPELD | SFKEELDKYF | KNHTSPDVDL | GDISGINASV | VNIQKEIDRL | NEVAKNLNES |
| QIA98583.1 | 1137 | VYDPLQPELD | SFKEELDKYF | KNHTSPDVDL | GDISGINASV | VNIQKEIDRL | NEVAKNLNES |
| QII57278.1 | 1137 | VYDPLQPELD | SFKEELDKYF | KNHTSPDVDL | GDISGINASV | VNIQKEIDRL | NEVAKNLNES |
| QIA20044.1 | 1137 | VYDPLQPELD | SFKEELDKYF | KNHTSPDVDL | GDISGINASV | VNIQKEIDRL | NEVAKNLNES |
| QIS60546.1 | 1137 | VYDPLQPELD | SFKEELDKYF | KNHTSPDVDL | GDISGINASV | VNIQKEIDRL | NEVAKNLNES |
| QIS60582.1 | 1137 | VYDPLQPELD | SFKEELDKYF | KNHTSPDVDL | GDISGINASV | VNIQKEIDRL | NEVAKNLNES |
| QIS61254.1 | 1137 | VYDPLQPELD | SFKEELDKYF | KNHTSPDVDL | GDISGINASV | VNIQKEIDRL | NEVAKNLNES |
| QIS61422.1 | 1137 | VYDPLQPELD | SFKEELDKYF | KNHTSPDVDL | GDISGINASV | VNIQKEIDRL | NEVAKNLNES |
| QIS60978.1 | 1137 | VYDPLQPELD | SFKEELDKYF | KNHTSPDVDL | GHISGINASV | VNIQKEIDRL | NEVAKNLNES |
| QIO04367.1 | 1137 | VYDPLQPELD | SFKEELDKYF | KNHTSPDVDL | GDISGINASV | VNIQKEIDRL | NEVAKNLNES |
| QIS30295.1 | 1137 | VYDPLQPELD | SFKEELDKYF | KNHTSPDVDL | GDISGINASV | VNIQKEIDRL | NEVAKNLNES |
| QIS30615.1 | 1137 | VYDPLQPELD | SFKEELDKYF | KNHTSPDVDL | GDISGINASV | VNIQKEIDRL | NEVAKNLNES |
| QIK50427.1 | 1137 | VYDPLQPELD | SFKEELDKYF | KNHTSPDVDL | GDISGINASV | VNIQKEIDRL | NEVAKNLNES |
| QIS30165.1 | 1137 | VYDPLQPELD | SFKEELDKYF | KNHTSPDVDL | GDISGINASV | VNIQKEIDRL | NEVAKNLNES |
| SARS-CoV | 1119 | VYDPLQPELD | SFKEELDKYF | KNHTSPDVDL | GDISGINASV | VNIQKEIDRL | NEVAKNLNES |

HR2

| | | | | | | | |
|---|---|---|---|---|---|---|---|
| YP_009724390 | 1197 | LIDLQELGKY | EQYIKWPWYI | WLGFIAGLIA | IVMVTIMLCC | MTSCCSCLKG | CCSCGSCCKF |
| QHS34546.1 | 1196 | LIDLQELGKY | EQYIKWPWYI | WLGFIAGLIA | IVMVTIMLCC | MTSCCSCLKG | CCSCGSCCKF |
| QIS60906.1 | 1197 | LIDLQELGKY | EQYIKWPWYI | WLGFIAGLIA | IVMVTIMLCC | MTSCCSCLKG | CCSCGSCCKF |
| QIS60930.1 | 1197 | LIDLQELGKY | EQYIKWPWYI | WLGFIAGLIA | IVMVTIMLCC | MTSCCSCLKG | CCSCGSCCKF |
| QIS61338.1 | 1197 | LIDLQELGKY | EQYIKWPWYI | WLGFIAGLIA | IVMVTIMLCC | MTSCCSCLKG | CCSCGSCCKF |
| QIS60489.1 | 1197 | LIDLQELGKY | EQYIKWPWYI | WLGFIAGLIA | IVMVTIMLCC | MTSCCSCLKG | CCSCGSCCKF |
| QIS30625.1 | 1197 | LIDLQELGKY | EQYIKWPWYI | WLGFIAGLIA | IVMVTIMLCC | MTSCCSCLKG | CCSCGSCCKF |
| QIS30335.1 | 1197 | LIDLQELGKY | EQYIKWPWYI | WLGFIAGLIA | IVMVTIMLCC | MTSCCSCLKG | CCSCGSCCKF |

```
QIC53204.1     1197 LIDLQELGKY EQYIKWPWYI WLGFIAGLIA IVMVTIMLCC MTSCCSCLKG CCSCGSCCKF
QHW06059.1     1197 LIDLQELGKY EQYIKWPWYI WLGFIAGLIA IVMVTIMLCC MTSCCSCLKG CCSCGSCCKF
QHZ00379.1     1197 LIDLQELGKY EQYIKWPWYI WLGFIAGLIA IVMVTIMLCC MTSCCSCLKG CCSCGSCCKF
QHR84449.1     1197 LIDLQELGKY EQYIKWPWYI WLGFIAGLIA IVMVTIMLCC MTSCCSCLKG CCSCGSCCKF
QIJ96493.1     1197 LIDLQELGKY EQYIKWPWYI WLGFIAGLIA IVMVTIMLCC MTSCCSCLKG CCSCGSCCKF
QIA98583.1     1197 LIDLQELGKY EQYIKWPWYI WLGFIAGLIA IVMVTIMLCC MTSCCSCLKG CCSCGSCCKF
QII57278.1     1197 LIDLQELGKY EQYIKWPWYI WLGFIAGLIA IVMVTIMLCC MTSCCSCLKG CCSCGSCCKF
QIA20044.1     1197 LIDLQELGKY EQYIKWPWYI WLGFIAGLIA IVMVTIMLCC MTSCCSCLKG CCSCGSCCKF
QIS60546.1     1197 LIDLQELGKY EQYIKWPWYI WLGFIAGLIA IVMVTIMLCC MTSCCSCLKG CCSCGSCCKF
QIS60582.1     1197 LIDLQELGKY EQYIKWPWYI WLGFIAGLIA IVMVTIMLCC MTSCCSCLKG CCSCGSCCKF
QIS61254.1     1197 LIDLQELGKY EQYIKWPWYI WLGFIAGLIA IVMVTIMLCC MTSCCSCLKG CCSCGSCCKF
QIS61422.1     1197 LIDLQELGKY EQYIKWPWYI WLGFIAGLIA IVMVTIMLCC MTSCCSCLKG CCSCGSCCKF
QIS60978.1     1197 LIDLQELGKY EQYIKWPWYI WLGFIAGLIA IVMVTIMLCC MTSCCSCLKG CCSCGSCCKF
QIO04367.1     1197 LIDLQELGKY EQYIKWPWYI WLGFIAGLIA IVMVTIMLCC MTSCCSCLKG CCSCGSCCKF
QIS30295.1     1197 LIDLQELGKY EQYIKWPWYI WLGFIAGLIA IVMVTIMLCC MTSCCSCLKG CCSCGSCCKF
QIS30615.1     1197 LIDLQELGKY EQYIKWPWYI WLGFIAGLIA IVMVTIMLCC MTSCCSCLKG CCSCGSCCKF
QIK50427.1     1197 LIDLQELGKY EQYIKWPWYI WLGFIAGLIA IVMVTIMLCC MTSCCSCLKG CCSCGSCCKF
QIS30165.1     1197 LIDLQELGKY EQYIKWPWYI WLGFIAGLIA IVMVTIMLCC MTSCCSCLKG CCSCGSCCKF
SARS-CoV       1179 LIDLQELGKY EQYIKWPWYV WLGFIAGLIA IVMVTILLCC MTSCCSCLKG ACSCGSCCKF
                     _______             __________________________________________________
                     HR2                               TM                        CT

YP_009724390   1257 DEDDSEPVLK GVKLHYT
QHS34546.1     1256 DEDDSEPVLK GVKLHYT
QIS60906.1     1257 DEDDSEPVLK GVKLHYT
QIS60930.1     1257 DEDDSEPVLK GVKLHYT
QIS61338.1     1257 DEDDSEPVLK GVKLHYT
QIS60489.1     1257 DEDDSEPVLK GVKLHYT
QIS30625.1     1257 DEDDSEPVLK GVKLHYT
QIS30335.1     1257 DEDDSEPVLK GVKLHYT
QIC53204.1     1257 DEDDSEPVLK GVKLHYT
QHW06059.1     1257 DEDDSEPVLK GVKLHYT
QHZ00379.1     1257 DEDDSEPVLK GVKLHYT
QHR84449.1     1257 DEDDSEPVLK GVKLHYT
QIJ96493.1     1257 DEDDSEPVLK GVKLHYT
QIA98583.1     1257 DEDDSEPVLK GVKLHYT
QII57278.1     1257 DEDDSEPVLK GVKLHYT
QIA20044.1     1257 DEDDSEPVLK GVKLHYT
QIS60546.1     1257 DEDDSEPVLK GVKLHYT
QIS60582.1     1257 DEHDSEPVLK GVKLHYT
QIS61254.1     1257 DEDDSEPVLK GVKLHYT
QIS61422.1     1257 DEDDSEPVLK GVKLHYT
QIS60978.1     1257 DEDDSEPVLK GVKLHYT
QIO04367.1     1257 DEDDSEPVLK GVKLHYT
QIS30295.1     1257 DEDDSEPVLK GVKLHYT
QIS30615.1     1257 DEDDSEPVLK GVKLHYT
QIK50427.1     1257 DEDDSEPVLK GVKLHYT
QIS30165.1     1257 DEDDSEPVLK GVKLHYT
SARS-CoV       1239 DEDDSEPVLK GVKLHYT
                    __________________
                           CT
```

Supplementary Fig 3. **Multiple sequence alignment of unique SARS-CoV-2 spike protein variants with SARS-CoV spike protein**. Identical regions are shaded throughout all the sequences.

Supplementary Table 1. **List of model quality assessment results of SARS-CoV-2 S protein variants in RAMPAGE server.**

| Variants | Number of residues | | |
|---|---|---|---|
| | **Favoured Region** | **Allowed Region** | **Outlier Region** |
| Template (pdb: 6VSB) | 892 (95.8%) | 38 (4.1%) | 1 (0.1%) |
| Y28N | 1025 (91.7%) | 74 (6.6%) | 19 (1.7%) |
| T29I | 1024 (91.6%) | 74 (6.6%) | 20 (1.8%) |
| H49Y | 1019 (91.1%) | 77 (6.9%) | 22 (2.0%) |
| L54F | 1023 ( 91.5%) | 73 ( 6.5%) | 22 ( 2.0%) |
| N74K | 1014 (90.7%) | 79 (7.1%) | 25 (2.2%) |
| E96D | 1021 ( 91.3%) | 73 ( 6.5%) | 24 ( 2.1%) |
| D111N | 1026 (91.8%) | 72 (6.4%) | 20 (1.8%) |
| F157L | 1028 (91.9%) | 71 (6.4%) | 19 (1.9%) |
| G181V | 1023 (91.5%) | 73 (6.5%) | 22 (2.0%) |
| S247R | 1024 (91.6%) | 74 (6.6%) | 20 (1.8%) |
| A348T | 1019 (91.1%) | 81 (7.2%) | 18 (1.6%) |
| R408I | 1022 ( 91.5%) | 77 ( 6.9%) | 18 ( 1.6%) |
| G476S | 1024 (91.6%) | 74 (6.6%) | 20 (1.8%) |
| V483A | 1030 (92.1%) | 74 (6.6%) | 14 (1.3%) |
| H519Q | 1023 (91.5%) | 71 (6.4%) | 24 (2.1%) |
| A520S | 1018 ( 91.1%) | 76 ( 6.8%) | 24 ( 2.1%) |
| D614G | 1020 ( 91.2%) | 75 ( 6.7%) | 23 ( 2.1%) |
| F797C | 1016 ( 90.9%) | 77 ( 6.9%) | 25 ( 2.2%) |
| A930V | 1022 ( 91.4%) | 73 ( 6.5%) | 23 ( 2.1%) |
| D936Y | 1027 (91.1%) | 71 (6.4%) | 20 (1.8%) |
| A1078V | 1023 ( 91.5%) | 72 ( 6.4%) | 23 ( 2.1%) |

Supplementary Table 2. **List of variations among 320 SARS-CoV-2 whole genomes.**

| Codon Variation | Mutation type | ORF |
|---|---|---|
| 8782C>T | Synonymous | orf1ab |
| 17747C>T | P5828L (17747C>T) | orf1ab |
| 17858A>G | Y5865C (17858A>G) | orf1ab |
| 18060C>T | Synonymous | orf1ab |
| 28144T>C | Synonymous | orf8 |
| 8602G>A | Synonymous | orf1ab |
| 26152G>A | G254R (26152G>A) | orf3a |
| 23120G>T | A520S | S |
| 2094C>T | S610L (2094C>T) | orf1ab |
| 19203T>C | Synonymous | orf1ab |
| 22984G>A | Synonymous | S |
| 27384T>C | Synonymous | orf6 |
| 241C>T | | 5' UTR |
| 1059C>T | T265I (1059C>T) | orf1ab |
| 3037C>T | Synonymous | orf1ab |
| 14408C>T | P4715L (14408C>T) | orf1ab |
| 23403A>G | D614G (23403A>G) | S |
| 25563G>T | Q57H (25563G>T) | orf3a |
| 29553G>A | Synonymous | orf10 |
| 34A>T | | 5' UTR |
| 35A>T | | 5' UTR |
| 36C>T | | 5' UTR |
| 37C>A | | 5' UTR |
| 15927T>G | Synonymous | orf1ab |
| 18401C>T | P6046L (18401C>T) | orf1ab |
| 24795C>T | A1078V (24795C>T) | S |
| 26936C>T | Synonymous | M |
| 11320T>C | Synonymous | orf1ab |
| 13845T>C | Synonymous | orf1ab |
| 16912G>T | V5550L (16912G>T) | orf1ab |
| 8768G>A | D2835N | orf1ab |
| 27525A>G | Synonymous | orf7a |
| 27722T>C | I110T (27722T>C) | orf7a |
| 8945A>G | N2894D (8945A>G) | orf1ab |
| 24022T>C | Synonymous | S |
| 28881G>A | R203K (28881G>A 28882G>A) | N |
| 28882G>A | R203K (28881G>A 28882G>A) | N |

| | | |
|---|---|---|
| 28883G>C | G204R (28883G>C) | N |
| 16887C>T | Synonymous | orf1ab |
| 25850G>A | C153Y (25850G>A) | orf3a |
| 21893G>A | D111N (21893G>A) | S |
| 27276C>T | Synonymous | orf6 |
| 1238C>T | H325Y (1238C>T) | orf1ab |
| 25494G>T | Synonymous | orf3a |
| 28924T>C | Synonymous | N |
| 16299T>C | Synonymous | orf1ab |
| 16329C>T | Synonymous | orf1ab |
| 23010T>C | V483A (23010T>C) | S |
| 4236A>G | N1324S (4236A>G) | orf1ab |
| 11083G>T | L3606F (11083G>T) | orf1ab |
| 14805C>T | Synonymous | orf1ab |
| 25655T>C | V88A (25655T>C) | orf3a |
| 26144G>T | G251V (26144G>T) | orf3a |
| 26028C>T | Synonymous | orf3a |
| 29866A>G | | 3' UTR |
| 29868G>C | | 3' UTR |
| 29869A>T | | 3' UTR |
| 8078C>T | P2605S (8078C>T) | orf1ab |
| 10319C>T | L3352F (10319C>T) | orf1ab |
| 15720C>T | Synonymous | orf1ab |
| 27964C>T | S24L (27964C>T) | orf8 |
| 17326C>T | P5688S (17326C>T) | orf1ab |
| 17884A>G | T5874A (17884A>G) | orf1ab |
| 19645G>T | V6461F (19645G>T) | orf1ab |
| 2232C>T | A656V (2232C>T) | orf1ab |
| 7518C>T | A2418V (7518C>T) | orf1ab |
| 23119T>A | H519Q (23119T>A) | S |
| 673C>T | Synonymous | orf1ab |
| 4683C>T | A1473V (4683C>T) | orf1ab |
| 29881A>C | | 3' UTR |
| 29888A>C | | 3' UTR |
| 25064G>C | D1168H (25064G>C) | S |
| 5A>T | | 5' UTR |
| 3593C>T | L1110F (3593C>T) | orf1ab |
| 27200A>G | Synonymous | M |
| 29862G>A | | 3' UTR |
| 29864G>A | | 3' UTR |
| 29867T>A | | 3' UTR |

| | | |
|---|---|---|
| 29868G>A | | 3' UTR |
| 29870C>A | | 3' UTR |
| 1440G>A | G392D (1440G>A) | orf1ab |
| 2891G>A | A876T (2891G>A) | orf1ab |
| 28854C>T | S194L (28854C>T) | N |
| 2416C>T | Synonymous | orf1ab |
| 26233G>T | | unknown |
| 12809C>T | Synonymous | orf1ab |
| 2041T>C | Synonymous | orf1ab |
| 11704C>T | Synonymous | orf1ab |
| 20256T>C | Synonymous | orf1ab |
| 29866A>T | | 3' UTR |
| 219G>T | | 5' UTR |
| 490T>A | D75E (490T>A) | orf1ab |
| 3177C>T | P971L (3177C>T) | orf1ab |
| 9113C>T | P2950S (9113C>T) | orf1ab |
| 18736T>C | F6158L (18736T>C) | orf1ab |
| 21648C>T | T29I (21648C>T) | S |
| 24034C>T | Synonymous | S |
| 26729T>C | Synonymous | M |
| 27635C>T | S81L (27635C>T) | orf7a |
| 28077G>C | V62L (28077G>C) | orf8 |
| 29567A>C | I4L (29567A>C) | orf10 |
| 29700A>G | | 3' UTR |
| 2676C>T | P804L (2676C>T) | orf1ab |
| 22606A>T | Synonymous | S |
| 28878G>A | S202N (28878G>A) | N |
| 29742G>A | | 3' UTR |
| 25337G>C | D1259H (25337G>C) | S |
| 29869A>C | | 3' UTR |
| 10323A>G | K3353R (10323A>G) | orf1ab |
| 79A>G | | 5' UTR |
| 23G>C | | 5' UTR |
| 84C>T | | 5' UTR |
| 85T>C | | 5' UTR |
| 90G>C | | 5' UTR |
| 29410T>C | Synonymous | N |
| 29557G>T | | unknown |
| 1A>T | | 5' UTR |
| 2T>G | | 5' UTR |
| 29880A>C | | 3' UTR |

| | | |
|---|---|---|
| 6310C>T | Synonymous | orf1ab |
| 27327G>T | K42N (27327G>T) | ORF6 |
| 28378G>T | Synonymous | N |
| 1401G>A | G379E (1401G>A) | orf1ab |
| 6693A>G | K2143R (6693A>G) | orf1ab |
| 14877C>T | Synonymous | orf1ab |
| 29188A>T | Synonymous | N |
| 619C>T | Synonymous | orf1ab |
| 14912A>G | N4883S (14912A>G) | orf1ab |
| 6040C>T | Synonymous | orf1ab |
| 12478G>A | M4071I (12478G>A) | orf1ab |
| 21910C>T | Synonymous | S |
| 28896C>G | A208G (28896C>G) | N |
| 29902A>C | | 3' UTR |
| 29903A>C | | 3' UTR |
| 9534C>A | T3090N (9534C>A) | orf1ab |
| 18877C>T | Synonymous | orf1ab |
| 28968G>C | S232T (28968G>C 28969C>T) | N |
| 28969C>T | S232T (28968G>C 28969C>T) | N |
| 2110C>T | Synonymous | orf1ab |
| 21575C>T | L5F (21575C>T) | S |
| 21850G>T | E96D (21850G>T) | S |
| 27804C>T | Synonymous | orf7b |
| 5000C>T | L1579F (5000C>T) | orf1ab |
| 29085C>T | T271I (29085C>T) | N |
| 5183C>T | P1640S (5183C>T) | orf1ab |
| 25691G>T | G100V (25691G>T) | orf3a |
| 28708C>T | Synonymous | N |
| 11916C>T | S3884L (11916C>T) | orf1ab |
| 55A>C | | 5' UTR |
| 1912C>T | Synonymous | orf1ab |
| 25452C>T | Synonymous | orf3a |
| 27246A>C | L15F (27246A>C) | orf6 |
| 27247C>T | L16S (27247C>T 27248T>C) | orf6 |
| 27248T>C | Synonymous | orf6 |
| 27255T>G | Synonymous | orf6 |
| 27340A>C | N47Q (27340A>C 27342T>G) | orf6 |
| 27342T>G | Synonymous | orf6 |
| 27344A>G | K48R (27344A>G) | orf6 |
| 8937C>T | A2891V (8937C>T) | orf1ab |
| 25523G>T | G44V (25523G>T) | orf3a |

| | | |
|---|---|---|
| 27869T>C | Synonymous | orf7b |
| 13748A>G | K4495R (13748A>G) | orf1ab |
| 1191C>T | P309L (1191C>T) | orf1ab |
| 21724G>C | L54F (21724G>C) | S |
| 12111G>A | S3949N (12111G>A) | orf1ab |
| 10818C>T | A3518V (10818C>T) | orf1ab |
| 14937C>T | Synonymous | orf1ab |
| 22988G>A | G476S (22988G>A) | S |
| 37C>T | | 5' UTR |
| 26433A>G | Synonymous | E |
| 28099C>T | S69L (28099C>T) | orf8 |
| 833T>C | F190L (833T>C) | orf1ab |
| 19137A>T | L6291F (19137A>T) | orf1ab |
| 8090C>T | L2609F (8090C>T) | orf1ab |
| 17247T>C | Synonymous | orf1ab |
| 565T>C | Synonymous | orf1ab |
| 17825C>T | T5854I (17825C>T) | orf1ab |
| 29573G>A | V6I (29573G>A) | orf10 |
| 29861G>A | | 3' UTR |
| 28280G>T | D3Y (28280G>T) | N |
| 29515T>A | Synonymous | N |
| 34A>C | | 5' UTR |
| 24368G>T | D936Y (24368G>T) | S |
| 6424A>C | E2053D (6424A>C) | orf1ab |
| 11042G>T | V3593F (11042G>T) | orf1ab |
| 22207T>C | Synonymous | S |
| 24694A>T | Synonymous | S |
| 28283A>G | N4D (28283A>G) | N |
| 20692C>T | P6810S (20692C>T) | orf1ab |
| 29845T>G | | 3' UTR |
| 29846T>A | | 3' UTR |
| 3T>C | | 5' UTR |
| 4A>T | | 5' UTR |
| 140C>T | | 5' UTR |
| 12213C>T | S3983F (12213C>T) | orf1ab |
| 19269C>T | Synonymous | orf1ab |
| 11674C>T | Synonymous | orf1ab |
| 28826C>T | R185C (28826C>T) | N |
| 197C>T | | 5' UTR |
| 198G>T | | 5' UTR |
| 200T>C | | 5' UTR |

| | | |
|---|---|---|
| 201T>A | | 5' UTR |
| 202T>G | | 5' UTR |
| 204G>C | | 5' UTR |
| 205T>G | | 5' UTR |
| 206C>G | | 5' UTR |
| 208G>T | | 5' UTR |
| 210G>A | | 5' UTR |
| 2632G>T | M789I (2632G>T) | orf1ab |
| 13225C>T | Synonymous | orf1ab |
| 14836A>G | I4858V (14836A>G) | orf1ab |
| 9180C>T | S2972F (9180C>T) | orf1ab |
| 29774C>T | | 3' UTR |
| 354G>A | G30D (354G>A) | orf1ab |
| 5716G>T | K1817N (5716G>T) | orf1ab |
| 19018C>T | P6252S (19018C>T) | orf1ab |
| 4642G>A | Synonymous | orf1ab |
| 27807C>T | Synonymous | orf7b |
| 14229G>T | K4655N (14229G>T) | orf1ab |
| 832C>T | Synonymous | orf1ab |
| 1545C>T | A427V (1545C>T) | orf1ab |
| 3992C>T | Synonymous | orf1ab |
| 27806G>A | Synonymous | orf7b |
| 28229T>G | H112Q (28229T>G) | orf8 |
| 14604A>G | Synonymous | orf1ab |
| 21911C>T | Synonymous | S |
| 19684G>T | V6474L (19684G>T) | orf1ab |
| 27999C>T | P36S (27999C>T) | orf8 |
| 19632T>A | N6456K (19632T>A) | orf1ab |
| 23989A>G | Synonymous | S |
| 17373C>T | Synonymous | orf1ab |
| 10572C>T | A3436V (10572C>T) | orf1ab |
| 11538T>C | I3758T (11538T>C) | orf1ab |
| 23242G>T | Synonymous | S |
| 28376G>A | A35T (28376G>A) | N |
| 6691T>C | Synonymous | orf1ab |
| 7843C>T | Synonymous | orf1ab |
| 13768A>G | M4502V (13768A>G) | orf1ab |
| 15869A>T | H5202L (15869A>T) | orf1ab |
| 17639C>T | S5792L (17639C>T) | orf1ab |
| 19542G>T | M6426I (19542G>T) | orf1ab |
| 28915C>T | Synonymous | N |

| | | |
|---|---|---|
| 3955G>T | K1230N (3955G>T) | orf1ab |
| 8681A>G | I2806V (8681A>G) | orf1ab |
| 22604G>A | A348T (22604G>A) | S |
| 150T>C | | 5' UTR |
| 153T>G | | 5' UTR |
| 29747G>A | | 3' UTR |
| 18433G>A | D6057N (18433G>A) | orf1ab |
| 29167C>T | Synonymous | N |
| 15888T>C | Synonymous | orf1ab |
| 313C>T | Synonymous | orf1ab |
| 1623T>C | I453T (1623T>C) | orf1ab |
| 3299T>C | Synonymous | orf1ab |
| 25156C>T | Synonymous | S |
| 29844A>G | | 3' UTR |
| 29847T>G | | 3' UTR |
| 29848T>G | | 3' UTR |
| 3099C>T | T945I (3099C>T) | orf1ab |
| 6636C>T | T2124I (6636C>T) | orf1ab |
| 11750C>T | L3829F (11750C>T) | orf1ab |
| 11956C>T | Synonymous | orf1ab |
| 16075G>T | D5271Y (16075G>T) | orf1ab |
| 18689T>A | F6142Y (18689T>A) | orf1ab |
| 20031C>A | Synonymous | orf1ab |
| 27046C>T | T175M (27046C>T) | M |
| 28409C>T | P46S (28409C>T) | N |
| 29736G>T | | 3' UTR |
| 3259G>T | Q998H (3259G>T) | orf1ab |
| 2269A>T | Synonymous | orf1ab |
| 29635C>T | Synonymous | orf10 |
| 9474C>T | A3070V (9474C>T) | orf1ab |
| 1385C>T | H374Y (1385C>T) | orf1ab |
| 29230C>T | Synonymous | N |
| 6026C>T | P1921S (6026C>T) | orf1ab |
| 12473C>T | Synonymous | orf1ab |
| 1469C>T | R402C (1469C>T) | orf1ab |
| 29254G>T | Synonymous | N |
| 10036C>T | Synonymous | orf1ab |
| 3T>G | | 5' UTR |
| 4A>G | | 5' UTR |
| 6A>T | | 5' UTR |
| 332G>T | V23F (332G>T) | orf1ab |

| | | |
|---|---|---|
| 5784C>T | T1840I (5784C>T) | orf1ab |
| 20823C>T | Synonymous | orf1ab |
| 2446T>C | Synonymous | orf1ab |
| 3411C>T | A1049V (3411C>T) | orf1ab |
| 5572G>T | M1769I (5572G>T) | orf1ab |
| 8092C>T | Synonymous | orf1ab |
| 26037C>T | Synonymous | orf3a |
| 27877G>T | C41F (27877G>T) | orf7b |
| 20281T>C | F6673L (20281T>C) | orf1ab |
| 254C>T | Synonymous | orf1ab |
| 29751G>C | | 3' UTR |
| 9157T>C | Synonymous | orf1ab |
| 22104G>T | G181V (22104G>T) | S |
| 28916G>A | G215S (28916G>A) | N |
| 10507C>T | Synonymous | orf1ab |
| 29871A>G | | 3' UTR |
| 6035A>G | S1924G (6035A>G) | orf1ab |
| 16467A>G | Synonymous | orf1ab |
| 21386C>T | S7041F (21386C>T) | orf1ab |
| 23185C>T | Synonymous | S |
| 29867T>C | | 3' UTR |
| 29871A>T | | 3' UTR |
| 29872A>C | | 3' UTR |
| 29873A>G | | 3' UTR |
| 29898A>G | | 3' UTR |
| 29902A>G | | 3' UTR |
| 29903A>G | | 3' UTR |
| 5052C>T | P1596L (5052C>T) | orf1ab |
| 17474C>T | T5737I (17474C>T) | orf1ab |
| 27226G>T | V9F (27226G>T) | orf6 |
| 1102C>T | Synonymous | orf1ab |
| 21147T>C | Synonymous | orf1ab |
| 22432C>T | Synonymous | S |
| 29837C>A | | 3' UTR |
| 514T>C | Synonymous | orf1ab |
| 17410C>T | R5716C (17410C>T) | orf1ab |
| 10232C>T | R3323C (10232C>T) | orf1ab |
| 221T>C | | 5' UTR |
| 27A>T | | 5' UTR |
| 18814C>T | Synonymous | orf1ab |
| 9477T>A | F3071Y (9477T>A) | orf1ab |

| | | |
|---|---|---|
| 25979G>T | G196V (25979G>T) | orf3a |
| 28657C>T | Synonymous | N |
| 28863C>T | S197L (28863C>T) | N |
| 4288G>T | E1341D (4288G>T) | orf1ab |
| 4307A>C | K1348Q (4307A>C) | orf1ab |
| 7479A>G | N2405S (7479A>G) | orf1ab |
| 11207G>C | A3648P (11207G>C) | orf1ab |
| 11233T>G | Synonymous | orf1ab |
| 12041G>C | D3926H (12041G>C) | orf1ab |
| 12160G>C | E3965D (12160G>C) | orf1ab |
| 12202G>C | K3979N (12202G>C) | orf1ab |
| 12208G>T | K3981N (12208G>T) | orf1ab |
| 12355G>C | Q4030H (12355G>C) | orf1ab |
| 12378G>A | R4038K (12378G>A) | orf1ab |
| 12464G>T | A4067S (12464G>T) | orf1ab |
| 12467G>T | A4068S (12467G>T) | orf1ab |
| 12491G>T | D4076Y (12491G>T) | orf1ab |
| 12514G>C | Synonymous | orf1ab |
| 12572G>T | D4103Y (12572G>T) | orf1ab |
| 12578G>T | D4105Y (12578G>T) | orf1ab |
| 12582G>T | S4106I (12582G>T) | orf1ab |
| 12600G>A | S4112N (12600G>A) | orf1ab |
| 12660G>C | R4132T (12660G>C) | orf1ab |
| 12685G>C | Q4140H (12685G>C) | orf1ab |
| 12773G>T | A4170S (12773G>T) | orf1ab |
| 12793G>T | K4176N (12793G>T) | orf1ab |
| 20980G>C | D6906H (20980G>C) | orf1ab |
| 21784T>A | N74K (21784T>A) | S |
| 9511T>C | Synonymous | orf1ab |
| 7G>A | | 5' UTR |
| 8G>C | | 5' UTR |
| 10T>A | | 5' UTR |
| 11161G>A | Synonymous | orf1ab |
| 18149G>T | Synonymous | orf1ab |
| 29865A>G | | 3' UTR |
| 10632C>T | A3456V (10632C>T) | orf1ab |
| 884C>T | R207C (884C>T) | orf1ab |
| 1348C>T | Synonymous | orf1ab |
| 1397G>A | V378I (1397G>A) | orf1ab |
| 9159C>T | P2965L (9159C>T) | orf1ab |
| 3373C>A | D1036E (3373C>A) | orf1ab |

| | | |
|---|---|---|
| 9561C>T | S3099L (9561C>T) | orf1ab |
| 15607T>C | Synonymous | orf1ab |
| 29095C>T | Synonymous | N |
| 17000C>T | T5579I (17000C>T) | orf1ab |
| 1548G>A | S428N (1548G>A) | orf1ab |
| 28792A>T | Synonymous | N |
| 8001A>C | D2579A (8001A>C) | orf1ab |
| 9534C>T | T3090I (9534C>T) | orf1ab |
| 7016G>A | G2251S (7016G>A | orf1ab |
| 21137A>G | K6958R (21137A>G) | orf1ab |
| 21316G>A | D7018N (21316G>A) | orf1ab |
| 24325A>G | Synonymous | S |
| 19065T>C | Synonymous | orf1ab |
| 22303T>G | S247R (22303T>G) | S |
| 7866G>T | G2534V (7866G>T) | orf1ab |
| 104T>A | | 5' UTR |
| 111T>C | | 5' UTR |
| 112T>G | | 5' UTR |
| 119C>G | | 5' UTR |
| 120T>C | | 5' UTR |
| 124G>A | | 5' UTR |
| 6996T>C | I2244T (6996T>C) | orf1ab |
| 614G>A | A117T (614G>A) | orf1ab |
| 5084A>G | I1607V (5084A>G) | orf1ab |
| 2091C>T | T609I (2091C>T) | orf1ab |
| 21707C>T | H49Y (21707C>T) | S |
| 2971G>T | M902I (2971G>T) | orf1ab |
| 6031C>T | Synonymous | orf1ab |
| 12115C>T | Synonymous | orf1ab |
| 15597T>C | Synonymous | orf1ab |
| 20936C>T | T6891M (20936C>T) | orf1ab |
| 22224C>G | S221W (22224C>G) | S |
| 25775G>T | W128L (25775G>T) | orf3a |
| 26354T>A | L37H (26354T>A) | E |
| 3518G>T | V1085F (3518G>T) | orf1ab |
| 17423A>G | Y5720C (17423A>G) | orf1ab |
| 75C>A | | 5' UTR |
| 11083G>C | L3606F (11083G>C) | orf1ab |
| 21644T>A | Y28N (21644T>A) | S |
| 9034A>G | H3076Y (9491C>T) | orf1ab |
| 9491C>T | Synonymous | orf1ab |

| | | |
|---|---|---|
| 186C>T | | 5' UTR |
| 2717G>A | G818S (2717G>A | orf1ab |
| 9274A>G | Synonymous | orf1ab |
| 13225C>G | F4321L (13226T>C) | orf1ab |
| 13226T>C | Synonymous | orf1ab |
| 17376A>G | Synonymous | orf1ab |
| 23952T>G | F797C (23952T>G) | S |
| 18603T>C | Synonymous | orf1ab |
| 18975T>A | Synonymous | orf1ab |
| 19175A>C | D6304A (19175A>C) | orf1ab |
| 27925C>T | T11I (27925C>T | orf8 |
| 9924C>T | A3220V (9924C>T) | orf1ab |
| 1A>C | | 5' UTR |
| 12534C>T | T4090I (12534C>T) | orf1ab |
| 13072C>T | Synonymous | orf1ab |
| 29873A>T | | 3' UTR |
| 29876A>G | | 3' UTR |
| 29877A>G | | 3' UTR |
| 29878A>T | | 3' UTR |
| 29880A>G | | 3' UTR |
| 29882A>G | | 3' UTR |
| 29886A>T | | 3' UTR |
| 29887A>T | | 3' UTR |
| 29889A>C | | 3' UTR |
| 29890A>T | | 3' UTR |
| 29891A>T | | 3' UTR |
| 29892A>G | | 3' UTR |
| 29893A>T | | 3' UTR |
| 29894A>C | | 3' UTR |
| 29897A>C | | 3' UTR |
| 29898A>T | | 3' UTR |
| 29901A>T | | 3' UTR |
| 29902A>T | | 3' UTR |
| 6819G>T | S2185I (6819G>T | orf1ab |
| 19610C>T | T6449I (19610C>T) | orf1ab |
| 29527G>A | Synonymous | N |
| 15324C>T | Synonymous | orf1ab |
| 29303C>T | P344S (29303C>T) | N |
| 29884A>G | | 3' UTR |
| 29885A>G | | 3' UTR |
| 29887A>G | | 3' UTR |

| | | |
|---|---|---|
| 654G>A | G130E (654G>A) | orf1ab |
| 4402T>C | Synonymous | orf1ab |
| 5062G>T | L1599F (5062G>T) | orf1ab |
| 29301A>T | D343V (29301A>T) | N |
| 1691A>G | I476V (1691A>G) | orf1ab |
| 6501C>T | P2079L (6501C>T) | orf1ab |
| 16877C>T | T5538I (16877C>T) | orf1ab |
| 24351C>T | A930V (24351C>T) | S |
| 2277T>C | I671T (2277T>C) | orf1ab |
| 6695C>T | P2144S (6695C>T) | orf1ab |
| 14657C>T | A4798V (14657C>T) | orf1ab |
| 22785G>T | R408I (22785G>T) | S |
| 3738C>T | P1158L (3738C>T) | orf1ab |
| 11410G>A | Synonymous | orf1ab |
| 26326C>T | Synonymous | E |
| 22033C>A | F157L (22033C>A) | S |
| 5845A>T | K1860N (5845A>T) | orf1ab |
| 3778A>G | Synonymous | orf1ab |
| 8388A>G | N2708S (8388A>G) | orf1ab |
| 8987T>A | F2908I (8987T>A) | orf1ab |

Supplementary Table 3. **List of SARS-CoV-2 Spike Protein Variants**.

| Variants | Frequency | Accession | SARS Residue[b] |
|---|---|---|---|
| L5F | 3 | QIS60906[a] | **L**5 |
| Y28N | 1 | QIA20044 | **Y**30 |
| T29I | 1 | QIS60546 | **T**31 |
| H49Y | 1 | QHW06059 | Y53 |
| L54F | 1 | QIS30295 | **L**58 |
| N74K | 1 | QIO04367 | Gap |
| E96D | 1 | QIS60930 | **E**93 |
| D111N | 1 | QIS61338 | N108 |
| Y145Del | 1 | QHS34546 | - |
| F157L | 1 | QII57278 | T150 |
| G181V | 1 | QIJ96493 | E174 |
| S221W | 1 | QHZ00379 | N214 |
| S247R | 1 | QHR84449 | Gap |
| A348T | 1 | QIS30335 | P335 |
| R408I | 1 | QHS34546 | **R**395 |
| G476S | 3 | QIS30625[a] | D463 |
| V483A | 4 | QIS30165[a] | P469 |
| H519Q | 1 | QIS61422 | N505 |
| A520S | 1 | QIS60489 | **A**506 |
| D614G | 71 | QIK50427[a] | **D**601 |
| F797C | 1 | QIC53204 | **F**779 |
| A930V | 1 | QIA98583 | **A**912 |
| D936Y | 1 | QIS30615 | E918 |
| A1078V | 1 | QIS61254 | **A**1060 |
| D1168H | 1 | QIS60978 | **D**1150 |
| D1259H | 1 | QIS60582 | **D**1241 |

[a]Supplementary Table 5 contain accession numbers of identical sequences of QIS60906, QIK50427, QIS30625, and QIS30165.

[b]Respective residue with position was counted from the MSA between spike protein of SARS-CoV and that of SARS-CoV-2.

Supplementary Table 4. **List of SARS-CoV-2 S protein sequences identical to SARS-CoV-2 reference S protein sequence (YP_009724390).**

| **YP_009724390** | | | | | |
|---|---|---|---|---|---|
| QHU79194 | QHU79204 | QIK50448 | QIQ49992 | QIS60399 | QIS61386 |
| QHZ00358 | QHZ87582 | QIA98554 | QIQ50132 | QIS60419 | QIS60858 |
| QHO62112 | QII87818 | QII57188 | QIQ50122 | QIS60469 | QIS61434 |
| QHQ71973 | QII87782 | QII57228 | QIQ49952 | QIS60429 | QIS60510 |
| QHU36824 | QII87794 | QIK02944 | QIQ50162 | QIS60276 | QIS60522 |
| QHU36834 | QII87806 | QIK02964 | QIQ50192 | QIS60439 | QIS60534 |
| QHZ00389 | QIC53213 | QIJ96463 | QIQ49862 | QIK02954 | QIS60570 |
| QHZ00399 | QIE07461 | QIJ96513 | QIQ50052 | QIH45033 | QIS60594 |
| QHU36864 | QIJ96473 | QII57248 | QIQ49842 | QHD43416 | QIS60678 |
| QHN73795 | QIE07451 | QII57298 | QIQ50182 | QIH45043 | QIS60690 |
| QHQ71963 | QIK50438 | QII57238 | QIQ49852 | QIH45053 | QIS60714 |
| QHW06049 | QIE07471 | QII57308 | QIQ49792 | QHO62107 | QIS61230 |
| QHQ82464 | QIJ96483 | QII57288 | QIQ49812 | QHR63260 | QIS61326 |
| QHO62877 | QHR63250 | QIG55994 | QIQ49822 | QHR63290 | QIS61350 |
| QHU36844 | QII57258 | QII57168 | QIQ50032 | QHW06039 | QIS60798 |
| QHU36854 | QIS30065 | QII57218 | QIQ50042 | QID21068 | QIS61398 |
| QIA98596 | QIS30018 | QII57268 | QIQ50062 | QHR63270 | QIS61410 |
| QIA98606 | QIS30030 | QII57338 | QIQ49912 | QHR63280 | QIS60966 |
| QIB84673 | QIS30255 | QIH45023 | QIQ49932 | QIQ08790 | QIS60990 |

| | | | | | |
|---|---|---|---|---|---|
| QHN73810 | QIS30325 | QID98794 | QIQ50002 | QIQ08830 | QIS61002 |
| QHZ87592 | QIS30345 | QID21058 | QIQ49892 | QIQ50012 | QIS61014 |
| QIE07481 | QIS30375 | QID21048 | QIQ68564 | QIQ50022 | QIS61026 |
| QIJ96503 | QIS30465 | QIS30545 | QIQ22760 | QIQ49962 | QIS61038 |
| QIJ96523 | QIS29994 | QIS30555 | QIS60726 | QIQ49982 | QIS61086 |
| QII57161 | QIS29982 | QIS30655 | QIS60738 | QIS60329 | QIS61098 |
| QIH55221 | QIS30006 | QIS30475 | QIS61266 | QIS60339 | QIS61146 |
| QIK50417 | QIS30175 | QIS30215 | QIS61302 | QIS60349 | QIS61158 |
| QII57178 | QIS30495 | QIS30042 | QIS61314 | QIS60449 | QIS61218 |
| QII57318 | QIS30515 | QIS30275 | QIS60786 | QIS60409 | QIS61482 |
| QII57208 | QIS30605 | QIS60479 | QIS61362 | QIS60459 | QIS61494 |
| QII57198 | QIS30485 | QIS60499 | QIS60359 | QIS60299 | QIS61506 |
| QII57328 | QIS30675 | QIS60309 | QIS60369 | QIS60319 | QIS61566 |
| QIQ68634 | QIQ68604 | QIQ68514 | QIS60379 | QIQ68684 | QIQ68554 |
| QHO60594 | QIQ68614 | QIQ68574 | QIS60389 | QIQ68544 | QIQ68644 |
| QIQ68494 | QIQ68654 | QIQ68584 | QIQ68674 | QIM47467 | QIQ68524 |
| QIQ68664 | QIQ68694 | QIQ68594 | QIQ68464 | QIQ68624 | |

Supplementary Table 5. **List of SARS-CoV-2 S protein sequences having similar variation.**

| Variant | Identical Sequences |
|---|---|
| **D614G** | QIK50427, QIS30235, QIS30245, QIS30265, QIS30365, QIS30385, QIS30435, QIS30445, QIS30155, QIS30305, QIS30095, QIS30195, QIS30415, QIS30315, QIS30405, QIS30535, QIS30585, QIS30665, QIS30125, QIS30075, QIS30085, QIS30285, QIS60288, QIQ49772, QIQ49972, QIQ50142, QIQ50112, QIQ50172, QIQ49872, QIQ50072, QIQ49762, QIQ49782, QIQ49802, QIQ50082, QIQ49942, QIS61290, QIS60870, QIS60618, QIS60630, QIS60642, QIS60666, QIS61242, QIS61278, QIS60762, QIS60810, QIS60834, QIS60894, QIS60918, QIS60942, QIS61050, QIS61074, QIS61122, QIS61134, QIS61182, QIS61194, QIS61206, QIS61446, QIS61518, QIS61542, QIS61554, QIQ68484, QIQ68474, QIQ68504, QIQ68534 |
| **L5F** | QIS60906, QIQ50152, QIQ49882 |
| **V483A** | QIS30165, QIS60774, QIS60882, QIS60954 |
| **G476S** | QIQ50152, QIQ49882, QIS30625 |
